# Cellular and molecular characterization of multiplex autism in human induced pluripotent stem cell-derived neurons

**DOI:** 10.1101/620807

**Authors:** Emily M.A. Lewis, Kesavan Meganathan, Dustin Baldridge, Paul Gontarz, Bo Zhang, Azad Bonni, John N. Constantino, Kristen L. Kroll

## Abstract

**Background:** Autism spectrum disorder (ASD) is a neurodevelopmental disorder with pronounced heritability in the general population. This is largely attributable to effects of polygenic susceptibility, with inherited liability exhibiting distinct sex differences in phenotypic expression. Attempts to model ASD in human cellular systems have principally involved rare *de novo* mutations associated with ASD phenocopies. However, by definition, these models are not representative of polygenic liability, which accounts for the vast share of population-attributable risk.

**Methods:** Here, we performed what is, to our knowledge, the first attempt to model multiplex autism using patient-derived induced pluripotent stem cells (iPSCs) in a family manifesting incremental degrees of phenotypic expression of inherited liability (absent, intermediate, severe). The family members share an inherited variant of unknown significance in *GPD2*, a gene that was previously associated with developmental disability but here is insufficient by itself to cause ASD. iPSCs from three first-degree relatives and an unrelated control were differentiated into both cortical excitatory (cExN) and cortical inhibitory (cIN) neurons, and cellular phenotyping and transcriptomic analysis were conducted.

**Results:** cExN neurospheres from the two affected individuals were reduced in size, compared to those derived from unaffected related and unrelated individuals. This reduction was, at least in part, due to increased apoptosis of cells from affected individuals upon initiation of cExN neural induction. Likewise, cIN neural progenitor cells from affected individuals exhibited increased apoptosis, compared to both unaffected individuals. Transcriptomic analysis of both cExN and cIN neural progenitor cells revealed distinct molecular signatures associated with affectation, including misregulation of suites of genes associated with neural development, neuronal function, and behavior, as well as altered expression of ASD risk-associated genes.

**Conclusions:** We have provided evidence of morphological, physiological, and transcriptomic signatures of polygenic liability to ASD from an analysis of cellular models derived from a multiplex autism family. ASD is commonly inherited on the basis of additive genetic liability. Therefore, identifying convergent cellular and molecular phenotypes resulting from polygenic and monogenic susceptibility may provide a critical bridge for determining which of the disparate effects of rare highly deleterious mutations might also apply to common autistic syndromes.

## Background

Autism spectrum disorder (ASD) is a neurodevelopmental disorder with a complex and poorly understood etiology (1-3). Behavioral and imaging studies have been valuable for defining deficits in affected individuals and characterizing alterations at the level of the brain. However, we are extremely limited in our ability to acquire or experimentally manipulate human brain tissue from living patients or post-mortem brain slices. This has hampered efforts to study cellular and molecular abnormalities that accompany ASD, both during and after fetal and post-natal development. Notably, both the relative integrity of brain structures in affected individuals and the diversity of ASD genetics suggest that convergent mechanisms that contribute to affectation in ASD may operate at the level of the cell (3, 4). These may be identifiable in experimental models derived from affected individuals. In particular, ASD appears to frequently involve abnormal development and/or function of two major classes of neurons in the cerebral cortex, glutamatergic excitatory projection neurons (cExNs) and GABAergic inhibitory interneurons (cINs) (3, 5, 6). *In vitro* differentiation-based models of these neuronal cell types can identify cellular and molecular deficits associated with ASD, and provide a tractable platform to screen for pharmacologic agents that can rescue these deficits.

In recent years, such cellular models of ASD have been generated either by deriving induced pluripotent stem cell lines (iPSCs) from affected individuals or by using CRISPR/Cas9-based gene editing to engineer ASD-associated mutations into wild type PSCs (7-19). Most of these studies have focused on syndromic forms of ASD, or on monogenic, *de novo* cases, where causality is attributed to mutation of a single ASD-linked gene (10, 11, 13-16). These forms of ASD are attractive for cellular modeling, as they streamline study design and reduce many potential confounding variables. Other studies have included individuals with an unknown genetic cause of ASD, but with subject selection based upon a shared phenotypic characteristic, such as macrocephaly (18-20). Together, these models have been informative, revealing both cellular and molecular alterations associated with affectation. These include shared and model-specific disruptions of gene expression in ASD-derived neurons, frequently involving altered expression of genes in key developmental signaling pathways, and genes that control cellular proliferation and growth (7, 8, 13-20). In addition, differences were observed in neural precursor cell (NPC) proliferation and differentiation (9, 19), neurogenesis, (9, 11, 18, 20), synaptogenesis (8, 10, 17, 18), or functional neuronal activity (7-9, 11, 16, 17). Altered expression of ASD genes, in which a mutation is linked to ASD causation or risk, is also frequently observed (7, 8, 13-16, 18-20).

These cellular modeling studies have revealed potential contributors to affectation and, in some cases, have identified targets amenable to pharmacological rescue *in vitro* (10, 18). However, they do not encompass the range of contributors to ASD burden in the general population: no single gene mutation accounts for more than 1% of overall ASD cases with predicted monogenic causality (21), while the majority of genetic risk appears to be polygenic or idiopathic (2, 22-28). Polygenic ASD risk can involve both common and rare variation in protein-coding and non-coding regions of the genome, which may act in a combinatorial manner (2, 27, 29, 30). Furthermore, ASD exhibits pronounced heritability in families (estimated at 50-90% (31)), none of which can be accounted for by *de novo* (germline) events. Even within each multiplex ASD family, there is often a considerable range in the extent of affectation among individuals, and in most multiplex autism families, a single causative gene mutation usually cannot be identified (31).

While ASD burden in the general population predominantly involves polygenic or idiopathic risk, heritability, and variable affectation (2, 21, 27, 29-31), these genetically complex forms of the disorder have been largely neglected in cellular modeling studies. Therefore, we deemed it important to determine whether cellular modeling of these complex but prevalent forms of ASD could also reveal affectation-related deficits. To test this, we focused on a multiplex ASD family with variable affectation among family members. We generated iPSC lines from three first degree relatives (a male and a female with differing degrees of affectation and their unaffected mother) as well as an unrelated unaffected female. We used these lines to perform differentiation into both cExN and cIN neural progenitors and/or neurons. Models from the affected individuals exhibited compromised cellular responses to differentiation cues and had disrupted gene expression profiles. This included altered expression of many ASD genes, genes with roles related to behavior, cognition, and learning, and genes involved in nervous system development and function, including cell adhesion molecules and ion channels.

This is, to our knowledge, the first cellular modeling study of multiplex ASD, including graded affectation among family members. We demonstrated that even genetically complex forms of ASD have discernable cellular and molecular abnormalities that track with affectation, some of which overlap those identified in prior modeling of syndromic, monogenic, and *de novo* forms of ASD. Therefore, this novel study design highlights the potential for cellular modeling to identify convergent hallmarks across the broad diversity and genetic complexity of pathways to affectation.

## Methods

### Phenotyping of the Multiplex Family

The nuclear family consisted of working professional non-consanguineous parents, whose first born child was a daughter with DSM-5 Autism Spectrum Disorder (ASD), Level I (requiring support, meeting DSM-5 criteria for Asperger Syndrome) who was very high functioning and ultimately attended college, followed by a pair of monozygotic twin boys with ASD, Level III— one more fluently verbal than the other but both severely impaired and requiring very substantial support (see below)—followed by a third son with very subtle autistic traits and predominantly affected by Attention Deficit Hyperactivity Disorder which improved substantially with stimulant medication treatment. Trio Exome sequencing (ES) of one of the twins and his parents revealed a variant of unknown significance (VUS) in *GPD2*, which was inherited by all of the children from the mother, who is of above average intelligence with no dysmorphism and no history of developmental problems. All pregnancies were uncomplicated, except for the post-natal hospital course of the twins.

The daughter was born at term with no complications or dysmorphia. Her language and motor development were typical and she was able to read at an early age. By age five she was reading at a fifth-grade level. She has been described as talented in writing and drawing. According to her parents, she exhibited social oddities from an early age, mainly in communication, and has somewhat intense/restricted interests in fantasy games. She has strong language abilities, and currently attends a four-year college, but at times uses odd phrases and the rhythm of her speech includes irregular pauses. She has described feeling alienated and “different”, and was the victim of bullying in middle school, with few close friends. In late adolescence, she developed major depressive disorder with moderate severity, which brought her to first psychiatric contact. She is cognizant of some degree of social awkwardness, which leads to feelings of anxiety and self-consciousness. The social anxiety inhibits her from activities such as eating in the cafeteria and pursuing job opportunities for which she is otherwise well-qualified. She has a history of becoming emotionally dysregulated and overwhelmed in times of stress, which has led to self-injurious behaviors. She has had ongoing struggles with depressive decompensation and suicidal ideation. She has above average intelligence but has struggled academically in college due to depression and anxiety. She is medically healthy with the exception of supraventricular tachycardia secondary to atrioventricular node reentry, which was treated with ablation and resulted in subsequent normalization of her electrocardiogram.

The twin boys were born at 35 weeks, had breathing problems at birth, and spent ten days in the newborn intensive care unit. Neither child has any dysmorphic features or congenital medical abnormalities, and brain imaging studies were negative. Likewise, neither child has a history of confirmed seizures, however, there are concerns for possible absence epilepsy. There is no history of abnormal neurological examination or macro-/microcephaly. Development of both siblings was delayed, but neither had appreciable regression. The more severely affected twin (designated as the affected proband, AP, and from whom the incuced pluripotent stem cell (iPSC) model of severe ASD affectation was acquired) began to exhibit delays in development by nine months of age. He was speaking single words at 14 months and was ultimately diagnosed with autism at 3.5 years old. Research confirmation of the diagnosis was obtained using the Autism Diagnostic Observation Schedule. Compared to his twin brother, he has had more perseverative interests on odd objects. Psychological testing at the age of nine revealed an IQ of 65 using the Leiter International Performance Scale. Now in late adolescence, he has the ability to engage in reciprocal and meaningful verbal exchanges, although his language is often echolalic and repetitive. He is socially motivated and develops superficial friendships with peers. Functionally, he is able to complete most self-care, dress himself, prepare food and feed himself, and count money. He participates in a vocational program at school and is able to complete rudimentary tasks assigned to him. His monozygotic twin was also diagnosed with ASD at 3.5 years old and is less severely affected, but still requires significant support. Selected clinical characteristics of the family members studied are summarized in Table 1.

**Table 1.** Selected clinical characteristics and mutational status of several individuals in the multiplex ASD pedigree.

| Clinical characteristic | AP | IS | UM |
| --- | --- | --- | --- |
| Age of ASD diagnostic confirmation | 3.5 years | 18 years | N/A |
| Social Responsiveness Scale-2 | 83T | 72T | 56T (spouse-report) |
| Depression and Anxiety | Yes | Yes | No |
| Seizure history | No | No | No |
| Developmental delay | Yes | No | No |
| Eye contact | Poor | Fair | Good |
| Repetitive behavior | Yes | No | No |
| Abnormal sensory sensitivities | Yes | No | No |
| IQ | 65 | 102 (Raven) | 108 (Raven) |
| Speech delay | Yes | No | No |
| ASD | Severe | Moderate | No |
| Intellectual Disability | Yes | No | No |
|  | GPD2 | GPD2 | GPD2 |
| Mutation location | chr2:157352686<br>(hg19) G>A | chr2:157352686<br>(hg19) G>A | chr2:157352686<br>(hg19) G>A |
|  | p.G78E, c.233G>A | p.G78E, c.233G>A | p.G78E, c.233G>A |

### Genotyping of the Multiplex Family

Trio ES was performed by GeneDx for the unaffected mother (UM), the AP, and the unaffected father. Briefly, sequencing was performed on an Illumina platform and reads were aligned to human genome build GRCh37/hg19 and analyzed using Xome Analyzer, as described (32). Mutation-specific testing was also performed by GeneDx on the intermediate phenotype sister (IS) and the third trait-affected brother (TB) to confirm presence of the identified *GPD2* variant in these individuals.

### iPSC generation

iPSC lines were generated by the Genome Engineering and iPSC Center at Washington University. Briefly, renal epithelial cells were isolated and cultured from fresh urine samples and were reprogrammed using a CytoTune-iPS 2.0 Sendai Reprogramming kit (Thermo Fisher Scientific), following the manufacturer’s instructions. iPSC clones were then picked, expanded, and characterized for pluripotency by immunocytochemistry (ICC) and RT-qPCR. At least three clonal iPSC lines were derived for each subject and two different clones were used for experimentation.

### iPSC maintenance and differentiation

iPSC lines were grown under feeder-free conditions on Matrigel (Corning) in mTeSR1 (STEMCELL Technologies). cExN and cIN differentiation of iPSCs was performed using previously described protocols (33). Briefly, for cExN differentiation, iPSCs were dissociated to single cells with Accutase (Life Technologies) and 40,000 cells were seeded in V-bottom 96-well non-adherent plates (Corning). Plates were spun at 200xg for five minutes to generate embryoid bodies (EBs) and were incubated in 5% CO_2_ at 37°C in cExN differentiation medium with 10µM Y-27632 (Tocris Biosciences). cExN differentiation medium components include Neurobasal-A (Life Technologies), 1X B-27 supplement (without Vitamin A) (Life Technologies), 10µM SB-431542 (Tocris Biosciences), 100nM LDN-193189 (Tocris Biosciences). On day four of differentiation, EBs were transferred from V-bottom plates to Poly-L-Ornithine-(20µg/ml) and laminin-(10µg/ml) coated plates. Media (without Y-27632) was replenished every other day, and on day 12 Neural Rosette Selection reagent (STEMCELL Technologies) was used to select neural progenitor cells (NPCs) from within neural rosettes, per the manufacturer’s instructions. cExN NPCs were grown as a monolayer using cExN differentiation media for up to 15 passages. cIN differentiation media contained the same components as cExN differentiation media, while also including 1µM Purmorphamine (Calbiochem) and 2µM XAV-939 (Tocris Biosciences). EBs were generated as described for cExNs. At day four of differentiation, the EBs were transferred to non-adherent plates and were placed on an orbital shaker (80 rpm) in an incubator with 5% CO_2_ at 37°C. The media was replenished every other day and, at day ten, EBs were transferred to Matrigel- and laminin-(5µg/ml) coated plates. Y-27632 was included in media until day eight of differentiation. On day 12 of differentiation, NPCs were dissociated with Accutase and maintained as a monolayer for up to 15 passages. For both cIN and cExN NPC growth analysis, an equal number of cells were seeded on Matrigel- and laminin-(5µg/ml) coated plates and total cells were counted four days later when the cells reached 70-80% confluence.

For differentiation of cExN NPCs into neurons for maturation, 40,000 cells per well were seeded in V-bottom 96-well non-adherent plates. Plates were spun at 200xg for five minutes and incubated in 5% CO_2_ at 37°C in maturation medium with Y-27632. Maturation medium components include Neurobasal-A, 1X B-27 supplement (without Vitamin A), 200µM cAMP (Sigma), 200µM Ascorbic acid (Sigma), and 20ng BDNF (Tocris Biosciences). After two days, EBs were transferred to Matrigel- and laminin-(5µg/ml) coated plates and media was replenished every other day (without Y-27632). On day 12 of neuronal differentiation and maturation, cells were dissociated with Accutase and seeded in an eight-well chamber for ICC.

For neurosphere size measurement analysis, p-values: \**P*<0.05, \*\**P*<0.01, \*\*\**P*<0.001 were determined by a two-tailed Student’s t-test.

### Sanger Sequencing

DNA was isolated from cell lines using the PureLink Genomic DNA Kit (Invitrogen). Primers were designed to amplify a 248 base pair region of *GPD2* flanking the identified point mutation (forward primer: AAGCAGCAGACTGCATTTCA, reverse primer: CACCATGGCACACACTTACC). Sanger sequencing was performed on this PCR amplified fragment using either the forward or reverse primer. CodonCode Aligner software was used to analyze sequencing results.

### Immunocytochemistry (ICC) and Immunoblotting

For ICC, cells were plated on eight-well chamber slides coated with Matrigel and laminin (5 µg/mL). After one day, cells were washed once with PBS without calcium and magnesium (PBS-Ca^2+^/Mg^2+^) and fixed in 4% Paraformaldehyde for 20 minutes, followed by washing with PBS + Ca^2+^/Mg^2+^. Cells were blocked with blocking buffer (10% donkey serum, 1% BSA, and 0.1% TritonX-100 in PBS + Ca^2+^/Mg^2+^) for at least one hour and incubated with primary antibodies overnight (Additional file 1: Table S1) in antibody dilution buffer (1% donkey serum, 1% BSA, and 0.1% TritonX-100 in PBS +Ca^2+^/Mg^2+^). After overnight incubation, cells were washed three times with wash buffer (0.1% Triton X-100 in PBS + Ca^2+^/Mg^2+^). Cells were incubated with corresponding secondary antibodies (Additional file 1: Table S1), along with DAPI (1mg/mL; ThermoFisher Scientific), diluted in antibody dilution buffer for one hour. Following secondary antibody incubation, cells were washed twice with wash buffer and once with PBS + Ca^2+^/Mg^2+^. Slides were mounted with Prolong Gold anti-fade agent (Life Technologies). Images were obtained using a spinning-disk confocal microscope (Quorum) with MetaMorph software and were processed using ImageJ. For immunoblotting, cell lysate was extracted and 30μg of protein was used per lane. Antibodies used are listed in Additional file 1: Table S1.

### FACS analysis

For each experiment, approximately one million cells were pelleted, washed with PBS – Ca^2+^/Mg^2+^, resuspended in PBS – Ca^2+^/Mg^2+^ and fixed by adding 70% ice-cold ethanol dropwise while vortexing. Cells were stained with 10ug/mL propidium iodide (Sigma) and 200ug/mL RNase A (Fisher Scientific) in FACS buffer (PBS – Ca^2+^/Mg^2+^, 0.2% BSA, 1mM EDTA). FACS was performed on single-cell suspensions and the cell cycle analysis function of FlowJo was used to analyze cell cycle composition for each sample, based on propidium iodide staining to detect DNA content in each cell. p-values: \**P*<0.05, \*\**P*<0.01, \*\*\**P*<0.001 were determined by a two-tailed Student’s t-test.

### RNA-Seq and RT-qPCR

Total RNA was collected from iPSC-derived day 12 cExN and cIN NPCs using the NucleoSpin RNA II kit (Takara) per the manufacturer’s instructions. RNA was quantified using a NanoDrop ND-1000 spectrophotometer (Thermo Scientific), and the integrity of RNA was confirmed with an Agilent Bioanalyzer 2100 to ensure a RIN value above eight. RNA-sequencing (RNA-seq) library preparation and Illumina sequencing were performed by the Genome Technology Access Center at Washington University. For RT-qPCR, 1µg total RNA was reverse transcribed using iScript Reverse Transcription Supermix (Bio-Rad). Equal quantities of cDNA were used as a template for RT-qPCR, using the Applied Biosystems Fast Real-Time quantitative PCR system. RPL30 mRNA levels were used as endogenous controls for normalization. p-values: \**P*<0.05, \*\**P*<0.01, \*\*\**P*<0.001 were determined by a two-tailed Student’s t-test.

### Bioinformatics and IPA analyses

RNA-seq data analysis was performed as described in (33) to curate differentially expressed gene (DEG) lists. To uncover the biological significance of DEGs, network analysis was performed with the data interpretation tool Ingenuity Pathway Analysis (IPA) (Qiagen). IPA’s Ingenuity Knowledge Base uses network-eligible DEGs to generate networks and to define connections between one or more networks. Based on the number of eligible DEGs, IPA defines network scores as inversely proportional to the probability of finding the network and defines significant networks (p≤0.001). Within each network, red symbols indicate upregulated genes and green symbols indicate downregulated genes, where the color intensity represents relative degree of differential expression.

## Results

### Phenotyping and genotyping of the multiplex family

The multiplex Autism Spectrum Disorder (ASD) pedigree selected for study (Fig. 1; Table 1) underwent clinical phenotyping and genotyping (see Methods). From this pedigree, the individuals selected for iPSC line derivation and modeling included the affected proband (AP), his sister, who has an intermediate phenotype (IS), and their unaffected mother (UM) (indicated in Fig. 1). As described above, a non-synonymous single nucleotide variant in the *GPD2* gene was identified in the UM and all of the children (chr2:157352686 (hg19) G>A, NM_001083112.2 c.233G>A, p.G78E). This variant is not present in the father. The variant is in exon three of the *GPD2* gene, within the region encoding the flavin adenine dinucleotide (FAD)-binding domain of the GPD2 protein (Additional file 2: Fig. S1A). The GeneDx interpretation of this variant states that it was not observed in approximately 6,500 individuals of European and African American ancestry in the NHLBI Exome Sequencing Project and that it is evolutionarily conserved. In addition, *in silico* analysis predicts that this variant is probably damaging to the protein structure and function. Overall, GeneDx designated this as a variant of uncertain significance (VUS) following the American College of Medical Genetics criteria. Subsequent mutation-specific testing and Sanger sequencing identified this same variant in the AP, IS and trait-affected brother (TB), while absent in an unrelated, unaffected control (UC) (Additional file 2: Fig. S1B). Given the differential affectation of members of this pedigree carrying this variant, it was apparent that this inherited VUS was insufficient to cause ASD by itself, but may have contributed to polygenic risk/liability in this family.

**Figure 1.**
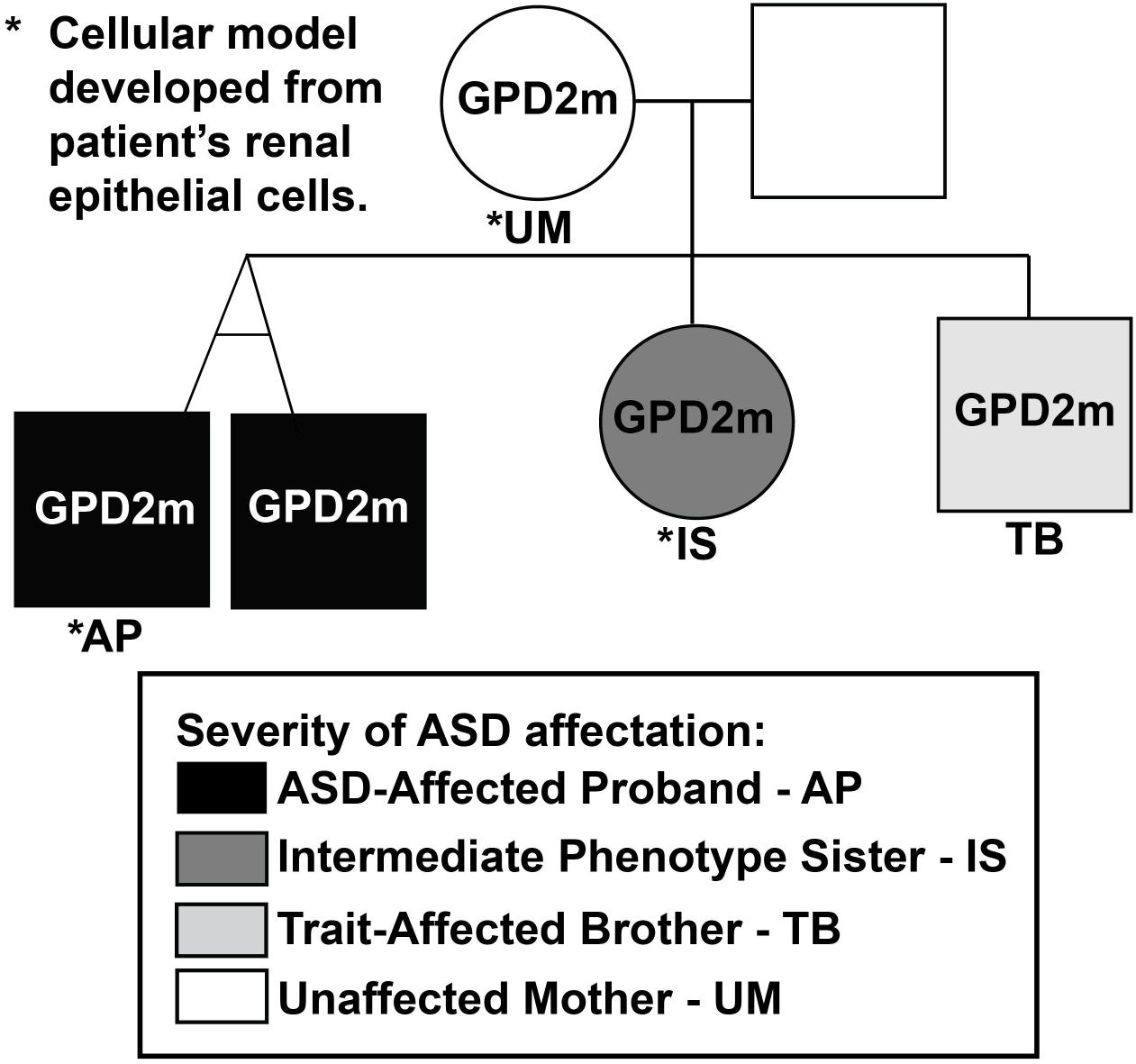
Pedigree from which samples were derived for this study. *GPD2* mutational status (GPD2m: indicates mutational status) and degree of ASD affectation are indicated. Black shading corresponds to the affected proband (AP) and his twin brother, dark grey to the intermediate phenotype sister (IS), light grey to the trait-affected brother (TB), and white to unaffected family members, including the unaffected mother (UM). * indicates that renal epithelial cells from these individuals were used to derive multiple, clonal iPSC lines.

### Generation of subject-derived iPSC models and directed differentiation into cortical excitatory neurons

Multiple clonal iPSC lines were derived from the UM, IS, and AP, and were characterized in parallel with a single, clonal iPSC line derived from the UC. When grown in stem cell maintenance media, there were no observable differences in expression of pluripotency markers between these iPSC lines, as assessed by RT-qPCR (Additional file 2: Fig. S1C) and immunocytochemistry (ICC) (Additional file 2: Fig. S1D). In addition, no differences in GPD2 protein levels were detected in iPSCs by ICC (Additional file 2: Fig. S1D) or Western blotting (Additional file 2: Fig. S1E). Finally, we used FACS analysis of propidium iodide (PI)-stained iPSCs to assess cell cycle progression and detected no observable differences between these iPSC lines, which had similar percentages of cells in each stage of the cell cycle (Additional file 2: Fig. S1F-G).

In the cortex, as a result of their abnormal development, imbalances in glutamatergic excitatory neurons (cExN) and GABAergic inhibitory interneurons (cINs) are thought to contribute to neurodevelopmental disorders including ASD (3, 5, 34). We therefore differentiated iPSCs derived from the UC, and from three family members (the UM, IS and AP), in parallel into either cExN or cIN neural progenitor cells (NPCs) and/or neurons, to determine if we observed any alterations in the *in vitro* development of either or both of these neural cell types. We performed 12 days of cExN differentiation (four days as embryoid bodies (EBs) in V-bottom plates, followed by eight days with the EBs plated for two-dimensional (2-D) culture; Fig. 2A). At all time points assessed during this differentiation, the IS and AP lines generated significantly smaller neurospheres than the UC and UM. The UM neurospheres were also slightly smaller than those of the UC (Fig. 2B-C).

**Figure 2.**
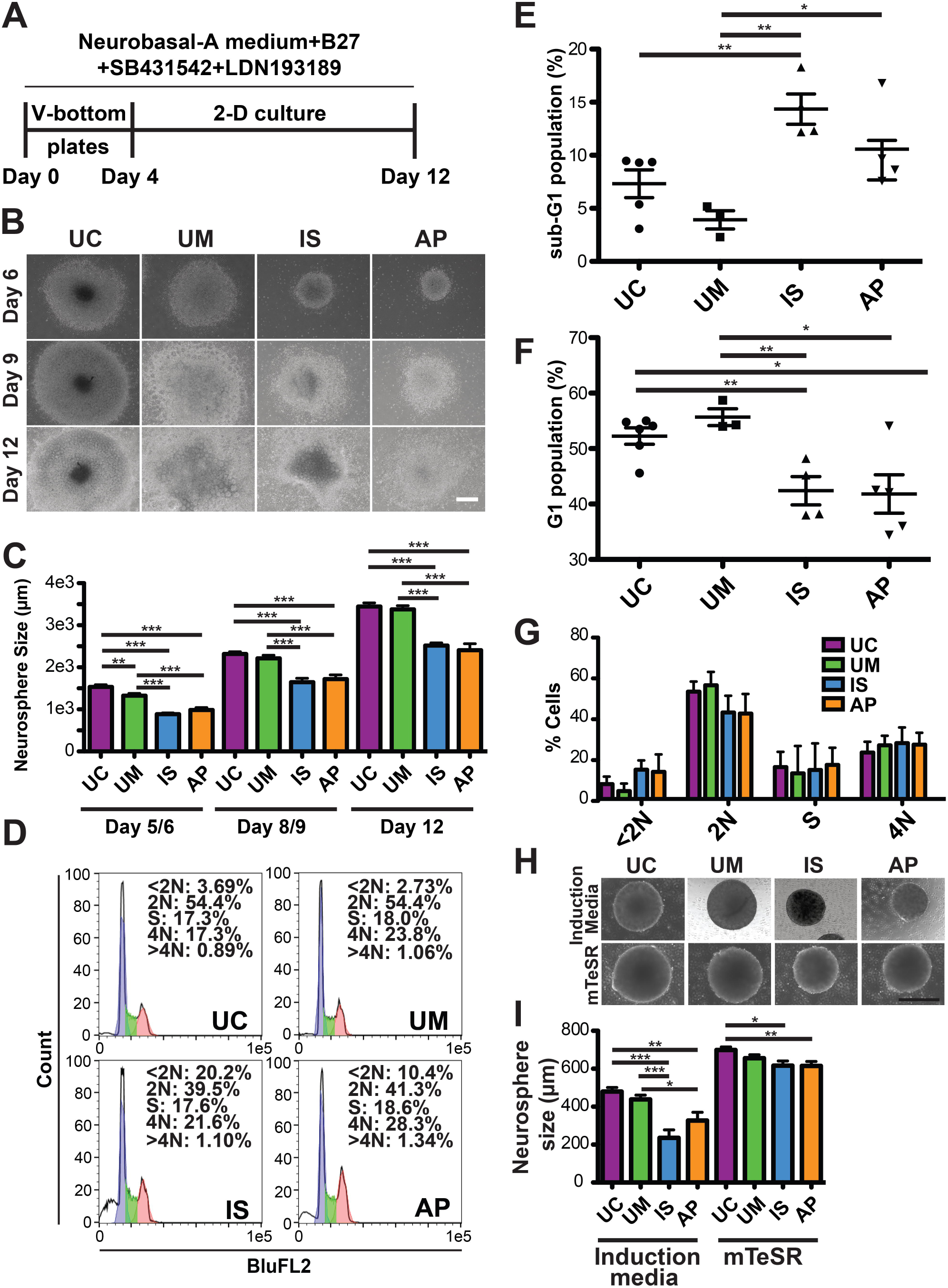
Characterization of iPSC lines during differentiation into cExN NPCs. **(A)** Differentiation scheme, including timeline and small molecules used. **(B-C)** iPSCs derived from an unrelated, unaffected control (UC), as well as the UM, IS, and AP were differentiated for 12 days to generate cExN NPCs. Neurosphere size at several time points is shown in (B) and quantified in (C) (mean±SEM; scale bar = 500µm; n=12 biological replicates, encompassing two different clonal lines from each subject, and one clonal line for the UC). **(D-G)** At day four of differentiation, cells were stained with propidium iodide and FACS analysis of DNA content was performed. (D) Representative FACS plots. In (E) <2N (sub-G1) and (F) 2N (G1) cells are quantified, with values shown for each replicate. (G) shows mean values for all cell cycle stages for each cell line (mean±SEM; n≥3 biological replicates for each subject, encompassing two different clonal lines from each subject, and one clonal line for the UC). **(H-I)** iPSCs were differentiated in either induction media or mTeSR stem cell media, and EB size was analyzed at day four of differentiation. Representative images are shown in (H) (scale bar = 500µm), with quantification in (I) (n=3 biological replicates from one clonal line for each subject). p-values: \**P*<0.05, \*\**P*<0.01, \*\*\**P*<0.001 were determined by a two-tailed Student’s t-test and all other pairwise comparisons had a non-signficant p-value (*P*≥0.05).

To identify whether an increase in apoptosis and/or a decrease in proliferation could be contributing to these differences in neurosphere size, we performed FACS analysis of PI-stained cells at day four of differentiation and found that the IS and AP neurospheres had a significantly higher fraction of sub-G1 (apoptotic) cells, compared to neurospheres derived from the UM and UC lines (<2N DNA content; Fig. 2D-E, G). There was a corresponding decrease in the percentage of cells in the G1 phase of the cell cycle in the IS and AP neurospheres (2N DNA content; Fig. 2D, F-G). However, neurospheres from all lines had similar percentages of cells in the S and G2/M phases of the cell cycle, suggesting that their cell cycle characteristics and rates of progression were otherwise similar (S phase and 4N DNA content; Fig. 2D, G). To determine whether induction of neural differentiation was a stressor that was contributing to this increase in apoptosis in the IS and AP line-derived neurospheres, we compared sphere size after culturing spheres from each line either in stem cell maintenance media (mTeSR) or in neural induction media. In general, sphere size was larger for all cell lines when kept in mTeSR media rather than neural induction media, while the differences in sphere size for the IS and AP versus the UC and UM was much less pronounced in mTeSR relative to differences seen under neural induction conditions (Fig. 2H-I). These data suggest that, by comparison with the UC and UM, the IS and AP lines have a slightly elevated propensity to undergo apoptosis upon dissociation and sphere formation, while this is exacerbated by induction of neural differentiation.

We next maintained these four lines as NPCs after neural rosette selection at day 12 and then subjected them to PI staining and FACS analysis. Unlike the results from earlier time points, the cExN NPCs showed no significant differences in cell cycle across the four lines (Additional file 2: Fig. S2A-B), nor in the rate of growth/death over the course of culture for four days (Additional file 2: Fig. S2C). However, morphological analysis by bright field imaging indicated a possible adhesion defect in the IS NPCs, as indicated by uneven growth on the cell culture plate surface (Additional file 2: Fig. S2D). At the NPC stage, GPD2 protein levels remained similar across the four lines, as was shown for iPSCs (Additional file 2: Fig. S2E).

Finally, to determine if the NPCs derived from the affected individuals exhibited an altered capacity to differentiate into cExN neurons, NPCs from the four lines were further differentiated for 12 days as shown (Additional file 2: Fig. S3A) and subjected to ICC. No apparent differences between the four lines were observed in the expression of NPC markers (PAX6, NESTIN, and SOX1) or markers of immature (TUJ1) and mature cExN neurons (VGLUT, MAP2) (Additional file 2: Fig. S3B). Furthermore, there were no observable differences between the lines in the fraction of cells expressing Ki-67, a marker of cell proliferation, or cleaved Caspase-3, a marker of apoptosis (Additional file 2: Fig. S3B).

### Differentiation of subject-derived iPSCs into cortical interneuron progenitors

We also characterized cellular phenotypes of these four lines during differentiation into cIN NPCs, to determine any differences between the development of this neural cell type in lines derived from affected versus unaffected individuals. The differentiation scheme to produce cIN NPCs is outlined in Figure 3A. On day five of differentiation in this scheme, neurospheres derived from the IS line were smaller than those of the UM. Conversely, the AP line-derived neurospheres were slightly larger than the UM line neurospheres (Fig. 3B-C).

**Figure 3.**
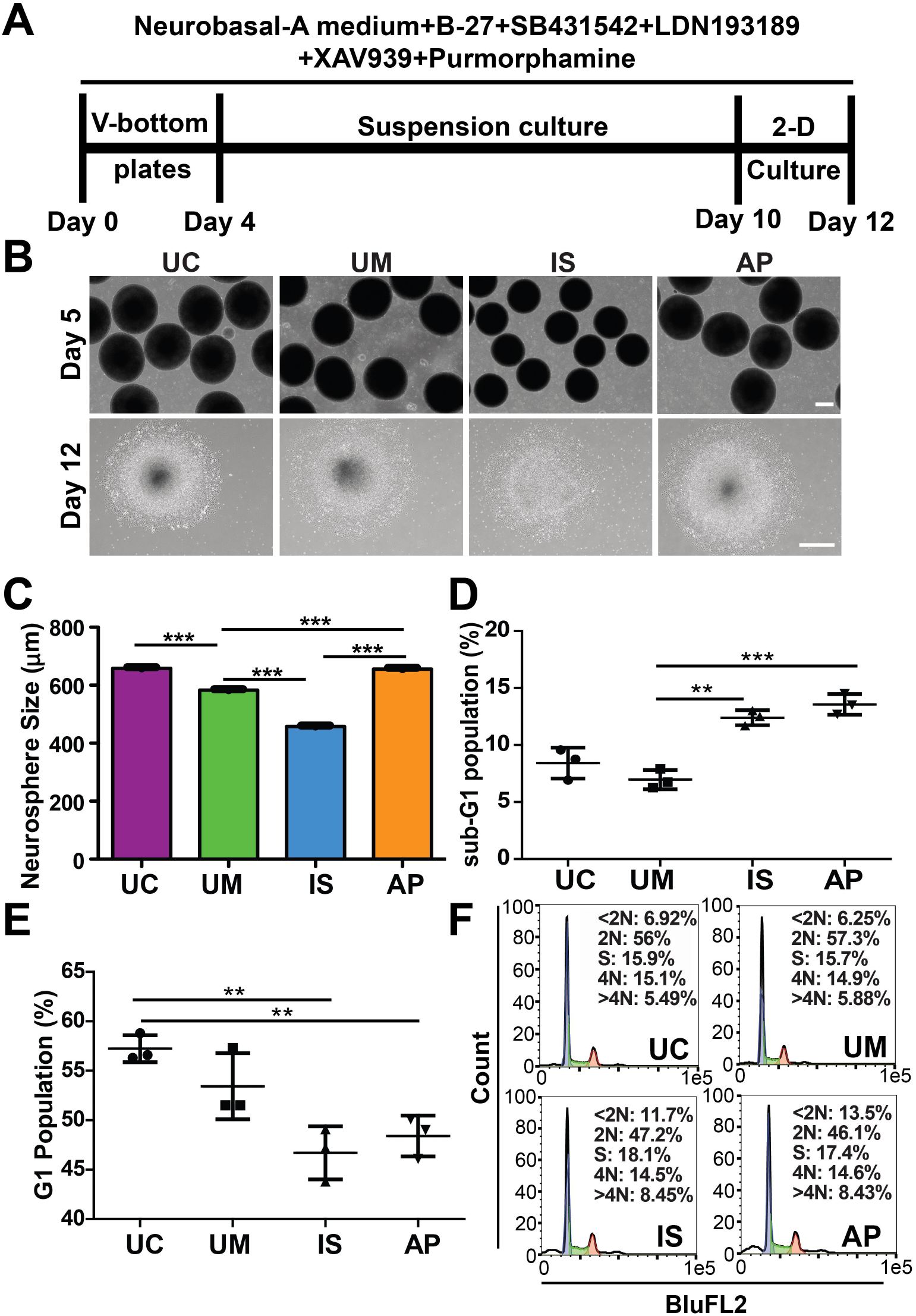
Characterization of iPSC lines during differentiation into cIN NPCs. **(A)** Differentiation scheme, including timeline and small molecules used. **(B-C)** iPSCs were differentiated for 12 days to obtain cIN NPCs, with differences in neurosphere size shown in (B) and quantified in (C) (mean±SEM; scale bar = 250µm; n=3 biological replicates from one clonal line for each subject). **(D-F)** NPCs were stained with propidium iodide and analyzed by FACS for DNA content. In D and E, respectively, <2N (sub-G1) and 2N (G1) cells were quantified, with values shown for each replicate (n=3 biological replicates from one clonal line for each subject). (F) shows representative FACS plots. p-values: \**P*<0.05, \*\**P*<0.01, \*\*\**P*<0.001 were determined by a two-tailed Student’s t-test and all other pairwise comparisons had a non-signficant p-value (*P*≥0.05).

After dissociation on day 12 of differentiation, we assessed the cell cycle of the cIN NPCs using FACS of PI-stained cells. The IS and AP cIN NPCs had an increased sub-G1 cell population, compared to the UC and UM NPCs, an indication of increased apoptosis in the cells from the affected individuals (Fig. 3D). Correspondingly, there was a decrease in the proportion of cells in the G1 phase of the cell cycle (Fig. 3E-F). However, no differences were observed in frequencies of cells in the S and G2 phases of the cell cycle between lines, suggesting that these lines had similar proliferation rates (Additional file 2: Fig. S2F). This result was supported by analysis of NPC cell counts after four days of growth, which revealed a significant reduction in the number of AP NPCs, as well as a slight reduction in the number of IS NPCs, compared to the UM NPCs (Additional file 2: Fig. S2G). These reductions in NPC number may result from the increased NPC apoptosis detected in our PI FACS analysis. The AP NPCs also exhibited altered morphology that could indicate impaired adhesion capacity relative to the control UM/UC lines, which could also contribute to the reduction in the number of AP NPCs persisting in the culture after four days of growth (Additional file 2: Fig. S2H).

### Transcriptomic differences in neural progenitor cells derived from affected individuals versus controls

To investigate which classes of genes could be differentially expressed in neural cells from the affected individuals, by comparison with the unaffected controls, we performed RNA-seq analysis on both cExN and cIN NPCs at day 12 of differentiation for all four subject-derived lines. Four biological replicates were analyzed for each sample type and were clustered by principal component analysis (PCA) of processed reads (Additional file 2: Fig. S4A-B). We defined genes that were significantly differentially expressed genes (DEGs) in pairwise comparisons of these four sample types for either cExN or cIN NPCs, selecting DEGs with a p-value of <0.05 and a fold difference between sample types of >2 (Additional file 3: Table S2). In a within-family comparison of the UM, IS, and AP samples, greater numbers of DEGs were obtained in the cIN NPC pairwise comparisons, versus the numbers of DEGs obtained for cExN NPC pairwise comparisons (Additional file 2: Fig. S4C-D). These data indicate that the cIN samples from the affected individuals (IS/AP) exhibit more transcriptomic differences from the UM control than the affected individual-derived cExN samples.

We focused first on identifying classes of genes that were differentially expressed in NPCs derived from the affected individuals, by comparison with unaffected controls. To do this, we defined the subset of DEGs that were similarly expressed in samples from both affected individuals (AP/IS) but that differed in expression by comparison with the unaffected mother (UM) sample. Relative expression is also shown for the UC, for a full cross-sample comparison. 452 and 437 DEGs for the cExN and cIN NPC samples met these criteria, respectively. Hierarchical clustering and visualization of the relative expression of these DEGs across the four sample types is shown for the cExN NPCs (Fig. 4A, Additional file 4: Table S3). We next used Ingenuity Pathway Analysis (IPA) to assess the potential biological significance of these genes. For the 452 DEGs in the cExN NPCs described above, the most significant function- and disease-related gene ontology (GO) terms included ‘behavior’, ‘neurological disease’, and ‘embryonic development’ (Fig. 4B). Network analysis using IPA revealed several interesting networks of DEGs related to these GO terms, including networks related to ‘locomotion’ (from DEGs within the ‘behavior’ GO term) and ‘behavior and developmental disorder’ (Fig. 4C-D). Within the ‘locomotion’ network, most genes were upregulated in the affected individuals compared to the controls, including genes relating to neural adhesion and ion channels (Fig. 4C and Additional file 5: Table S4). Genes with known roles in NPCs or neurons, as well as stress-related genes were present in the larger ‘behavior and developmental disorder’ network (Fig. 4D and Additional file 5: Table S4). Interestingly, another network comprising genes from the GO term ‘neurological disease’ is related to ‘inflammation of central nervous system’ (Additional file 2: Fig. S5A and Additional file 5: Table S4).

**Figure 4.**
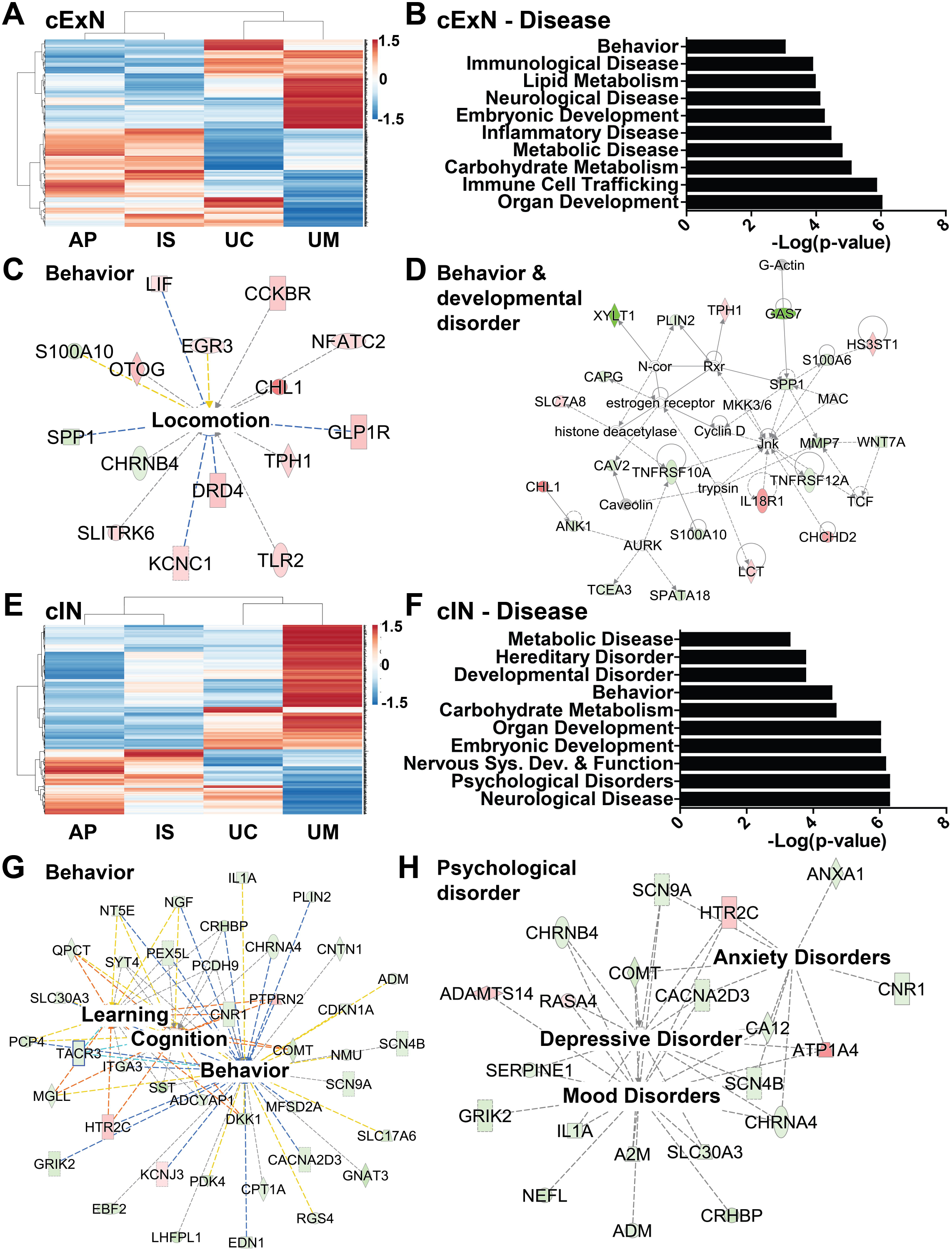
Transcriptomic analysis of genes with differential expression in affected subject-derived NPCs, relative to unaffected controls. **(A)** Hierarchical clustering of RNA-seq data from the cExN NPCs identified differentially expressed genes (DEGs) with shared expression in the AP and IS that differed from that in the UM. Relative expression is also shown for the UC, for a full cross-sample comparison (p<0.05, fold-change >2; n=4 biological replicates from one clonal line for each subject). **(B-D)** Ingenuity Pathway Analysis (IPA) of these cExN NPC DEGs defined disease and function-associated gene ontology (GO) terms and identified gene networks associated with the (C) ‘behavior’ (D) and ‘behavior and developmental disorder’ GO terms. **(E)** Hierarchical clustering of RNA-seq data for the cIN NPCs identified DEGs with shared expression in the AP and IS, that differed from that in the UM. Relative expression is also shown for the UC, for a full cross-sample comparison (p<0.05, fold-change >2; n=4 biological replicates from one clonal line for each subject). **(F-H)** IPA analysis of these cIN NPC DEGs defined (F) disease- and function-associated GO terms and identified gene networks associated with (G) ‘behavior’ and (H) ‘psychological disorder’. Within each network, red symbols indicate upregulated genes and green symbols indicate downregulated genes, while color intensity indicates relative degree of differential expression.

IPA analysis of the cIN DEGs also revealed several interesting classes of genes that were differentially expressed between the affected participants and unaffected control NPC-derived samples. Hierarchical clustering and visualization of the relative expression of DEGs across the four sample types for the cIN NPCs is shown in Fig. 4E (Additional file 4: Table S3). The top GO terms included ‘developmental disorder’, ‘behavior’, ‘nervous system development and function’, ‘psychological disorders’, and ‘neurological disease’ (Fig. 4F). Within the term ‘behavior’, a network that includes ‘learning’-, ‘cognition’-, and ‘behavior’-related genes was identified (Fig. 4G, Additional file 5: Table S4). The network related to the GO term ‘psychological disorder’ includes genes related to ‘anxiety disorders’, ‘mood disorders’, and ‘depressive disorder’ (Fig. 4H). The ‘nervous system development and function’ network includes genes involved in the ‘quantity of neurons’ and ‘quantity of synapse’, as well as cell adhesion genes (Additional file 2: Fig. S5B and Additional file 5: Table S4). Finally, a ‘neurological’ network included a number of genes also present in the other networks (Additional file 2: Fig. S5C). Taken together, this analysis of DEGs in both cExN and cIN NPCs shows evidence of altered expression of a number of neurological and psychological disease-relevant gene classes in the AP- and IS-derived lines, relative to lines derived from the UM and/or UC.

### Within-family comparison identifies a transcriptome signature specific to neural progenitor cells derived from the ASD-affected proband

As differences in genetic background can confound differential gene expression analysis (35), we also performed a pairwise, within-family data comparison of DEGs that distinguish the UM-, IS-, and AP-derived samples, focusing on DEGs specific to the AP that could contribute to the greater degree of affectation observed. Using pairwise comparisons of DEGs, we defined 190 genes which were uniquely differentially expressed in cExN NPCs from the AP (Fig. 5A). The top GO terms associated with these AP-specific DEGs included ‘psychological disorders’, ‘behavior’, ‘nervous system development and function’, ‘developmental disorder’, and ‘neurological disease’ terms (Fig. 5B). Within the ‘behavior’ term, network analysis showed genes related to ‘memory’ and ‘learning’ to be dysregulated (Fig. 5C and Additional file 5: Table S4). Within the ‘nervous system development and function’ term, a network of dysregulated genes related to ‘differentiation of neurons’ was identified (Fig. 5D, Table S4).

**Figure 5.**
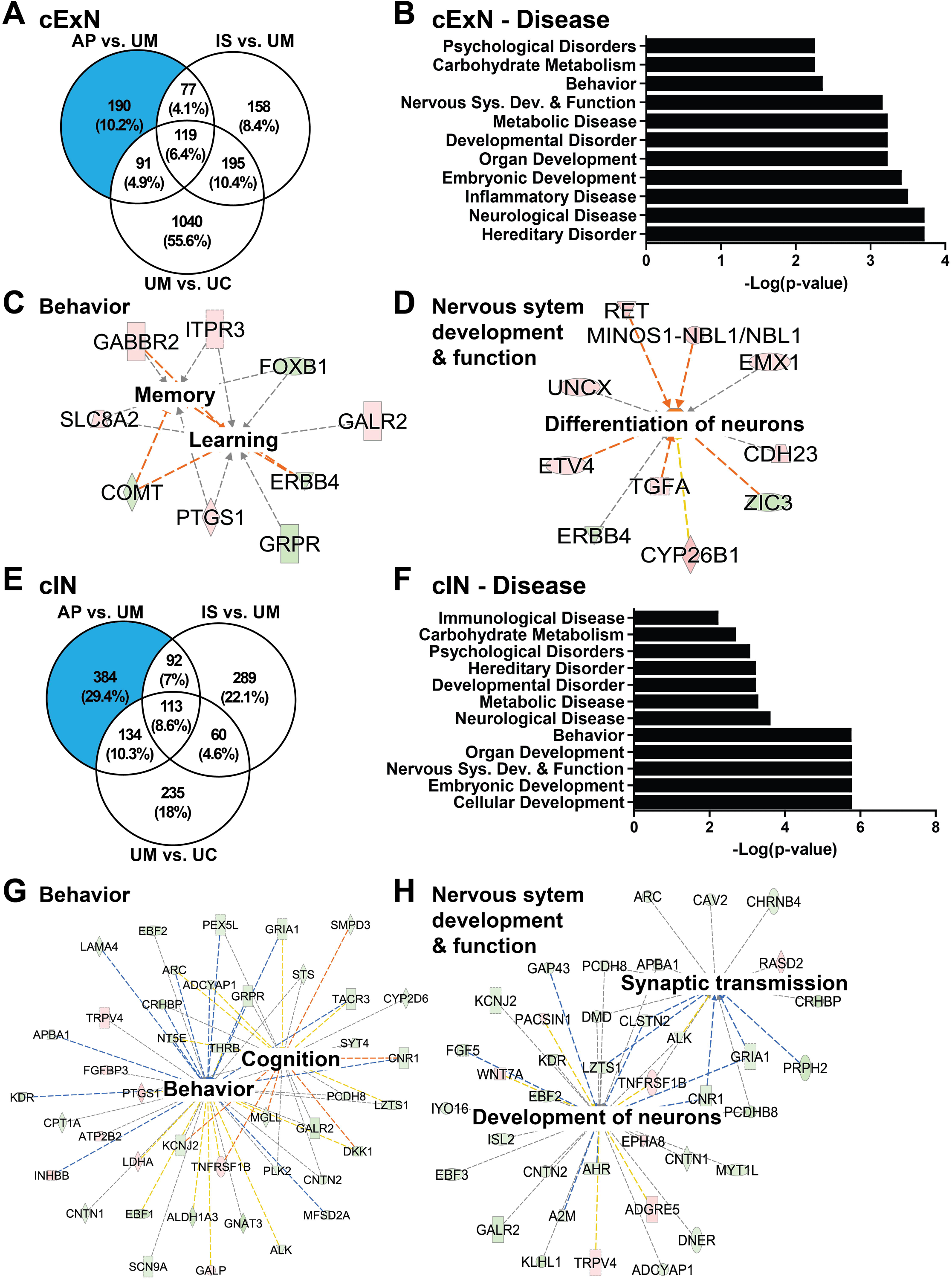
Within-family analysis of transcriptomic signatures specific to the affected proband-derived samples. **(A)** Venn diagram for the cExN NPCs, showing the DEGs from pairwise comparisons of different samples, including numbers of overlapping DEGs. The blue shaded portion of the Venn diagram indicates DEGs unique to the AP, not shared by the IS or UM. **(B-D)** Ingenuity Pathway Analysis (IPA) of the AP-unique DEGs in cExN NPCs defined class and function-associated GO terms (B) and identified gene networks associated with ‘behavior’ (C) and ‘nervous system development and function’ (D). **(E-H)** IPA analysis of the AP-unique DEGs in cIN NPCs determined (F) class and function-associated GO terms and identified gene networks associated with (G) ‘behavior’ and (H) ‘nervous system development and function’. Within each network, red symbols indicate upregulated genes and green symbols indicate downregulated genes, where the color intensity represents relative degree of differential expression.

Similar analysis was performed on the cIN samples, revealing 384 DEGs unique to the AP samples in the within-family comparison (Fig. 5E). IPA analysis identified classes of DEGs related to the GO terms ‘psychological disorders’, ‘developmental disorder’, ‘neurological disease’, ‘behavior’, and ‘nervous system development and function’ (Fig. 5F). Within the ‘behavior’ disease term, a network of genes related to ‘behavior’ and ‘cognition’ was identified (Fig. 5G and Additional file 5: Table S4). Within the ‘nervous system development and function’ term, a network of genes related to ‘development of neurons’ and ‘synaptic transmission’ had altered expression in the AP versus the IS/UM-derived samples (Fig. 5H and Additional file 5: Table S4). Together, this analysis identifies AP-unique DEGs in both cExN and cIN NPCs, many of which are broadly related to neural development, as well as to specific aspects of ASD, such as behavioral alterations. These gene expression changes therefore correlate with and may contribute to, the severity of affectation in the AP.

### Comparison of differentially expressed genes with ASD-associated genes and validation

The Simons Foundation Autism Research Initiative (SFARI) (36) maintains a database of genes that are mutated to cause, or contribute to, ASD risk. We compared our DEGs to these ASD genes, to assess whether their dysregulated expression could contribute to affectation in these individuals. Of 452 DEGs in the cExN differentiation scheme that had similar expression in the AP and IS that differed from that seen in the UM, 30 (6.6%) were ASD genes (Fig. 6A). For the corresponding cIN NPC comparison, 46 of 437 DEGs (10.5%) were ASD genes in the SFARI Gene database (Fig. 6B). Based upon the 1019 genes present in the SFARI Gene database (36) and the total of 27,731 genes with >0.1 RPKM average expression across all cExN and cIN samples, the number of AP- and IS-specific DEGs that are AD genes is significantly greater than would be expected by chance (hypergeometric distribution, p=9.35×10^−5^ and 1.95×10^−12^ for cExN and cIN data, respectively). Therefore, it is possible that misregulated expression and consequently function of ASD genes contributed to disruption of neural development and/or continues to contribute to altered neurological function in the IS and AP.

**Figure 6.**
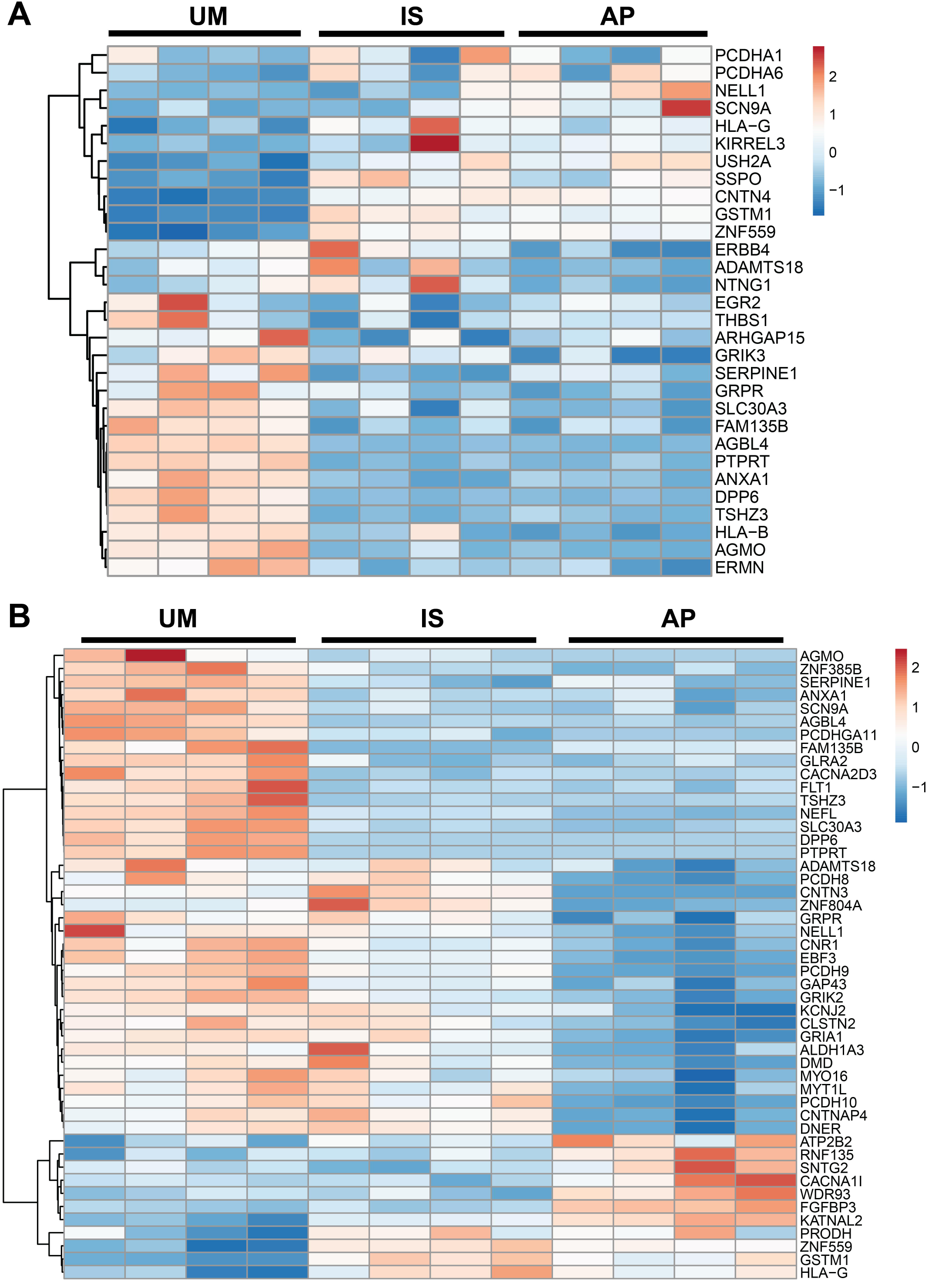
Hierarchical clustering of DEGs that are also ASD genes in the SFARI autism gene database. **(A)** Relative gene expression for the cExN NPC samples. **(B)** Relative gene expression for the cIN NPC samples. Data from four biological replicates are shown for each sample type.

We validated the differential expression of a subset of the DEGs described above by RT-qPCR analysis, isolating RNA from NPCs derived from a second set of iPSC clones that differed from those used for the RNA-seq experiments. Expression changes of DEGs selected from the cExN NPC RNA-seq data (Fig. 7A) were robustly recapitulated in these experiments (Fig. 7B). Genes from the ‘behavior and developmental disorder’, from other identified networks, and genes involved in neurodevelopment were evaluated (Additional file 5: Table S4). We also derived cIN NPC RNA from a second set of iPSC clones and validated the corresponding RNA-seq data for a subset of the DEGs (Fig. 7C). Differential expression was assessed for SFARI ASD genes (36), and for genes encoding transcription factors, ion channels, and cell adhesion molecules (Fig. 7D-E and Additional file 5: Table S4). In addition, we validated a subset of DEGs in cINs which also had differential expression in cExN NPCs (Fig. 7E). Together, these analyses revealed that, relative to unaffected individuals, samples from affected individuals exhibited altered expression of classes of genes involved in behavior, learning, cognition, mood disorders, and neurodevelopment, including perturbed ASD gene expression, suggesting that these differences could contribute to aberrant neural development or function in the affected individuals.

**Figure 7.**
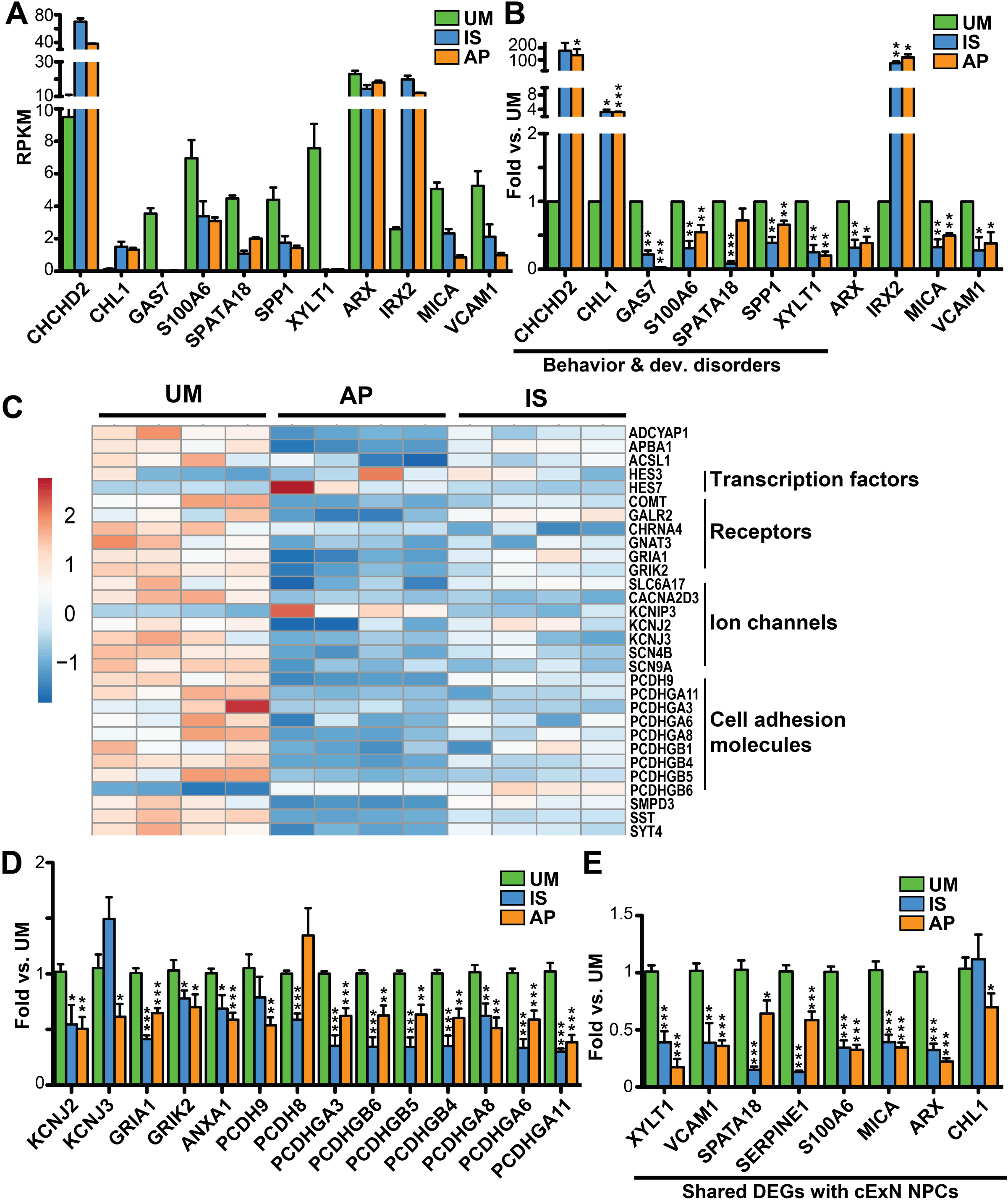
Validation of DEGs of interest identified from RNA-seq experiments by RT-qPCR. Genes tested are related to behavior and developmental disorders, adhesion, and ion channels. **(A-B)** Comparison of relative gene expression in cExN NPCs for the UM, IS, and AP by (A) RNA-seq and (B) RT-qPCR, including expression analysis of genes related to ‘behavior and developmental disorders’. **(C-E)** Comparison of gene expression between the UM, IS and AP for the cIN NPCs by (C) RNA-seq and by (D-E) RT-qPCR, both for genes that were (D) differentially expressed only in the cIN NPCs and (E) for genes that were differentially expressed in both cExN and cIN NPCs. p-values: \**P*<0.05, \*\**P*<0.01, \*\*\**P*<0.001 were determined by a two-tailed Student’s t-test and all other pairwise comparisons had a non-signficant p-value (*P*≥0.05). RT-qPCR data shown includes n≥3 biological replicates from one clonal line per subject, where samples were generated for each subject by using a second clonal iPSC line that differed from the line used for RNA-seq analysis.

## Discussion

In recent years, the genetic structure of Autism Spectrum Disorder (ASD) risk in the general population has been clarified. This work has confirmed that while, in some cases, deleterious, single gene variants are significant contributors to ASD, the vast proportion of population attributable risk is polygenic (2, 37). Furthermore, this risk is highly heritable, and individuals within a multiplex family typically exhibit variable degrees of affectation (31). Here, we modeled cellular and molecular correlates of ASD within one such multiplex family, performing cortical neural differentiation of iPSCs derived from several family members with differential affectation. In this family, both polygenic liability and a shared variant of unknown significance (VUS) may contribute to risk. In cells derived from the affected individuals, we identified compromised responses to differentiation cues and altered gene expression profiles during iPSC differentiation into cortical excitatory (cExN) and inhibitory (cINs) neurons, compared to related and unrelated unaffected controls. This work demonstrates that iPSC-based modeling can be used to characterize these more genetically complex but prevalent forms of ASD, in addition to modeling simplex and monogenic forms, which have been the focus of most studies to date. Moreover, these data provide information on physiologic and transcriptomic signatures of multiplex autism, with which cellular models derived from other families and other combinations of inherited susceptibility factors can be compared in future work.

Our phenotypic analysis of these four iPSC-based models of cortical neural development included assays conducted in the stem cells, during neural specification, in the proliferating NPCs, and during neuronal differentiation. During cExN NPC specification and during cIN NPC propagation, models from both affected individuals exhibited elevated fractions of cells with sub-G1 DNA content, relative to control-derived models. These data suggest that models derived from the affected individuals are less resistant to stressors, such as induction of differentiation, with these stressors increasing the propensity for cells to undergo apoptosis. While the molecular trigger for the induction of apoptosis here is unclear, expression of stress and apoptosis-related genes, such as *CHCHD2, ANXA1, and SPATA18* are dysregulated in these models (38-42). These findings are reminiscent of some observations made in prior work, in which schizophrenia subject-derived iPSCs exhibited reduced neurosphere size (43) and increased apoptosis was observed in Williams-Syndrome iPSC-derived NPCs (44). Interestingly, few studies report cellular alterations observed prior to the NPC stage, often focusing predominantly on phenotypes seen in NPCs and mature neurons (7-19). While this may reflect a lack of earlier phenotypic changes in some models, our findings highlight the importance of tracking neurodevelopmental alterations from their earliest onset. A recent report underscores the value of using cellular modeling approaches that aim to recapitulate some aspects of *in vivo* neurodevelopment (20). This study found that direct conversion of iPSCs into neurons masked ASD-associated cellular phenotypes, which were observable during directed differentiation of iPSCs (20).

In our study, transcriptomic analysis of neural progenitor cells revealed dysregulated expression in affected individuals compared to controls of gene networks related to behavior, psychological disorders, and neuronal development and disease. Genes encoding transcription factors were among the neurodevelopment-related genes with reduced expression in both affected individuals. For example, ARX is required for normal telencephalic development and is associated with syndromic autism and other neurodevelopmental disorders (45), while EMX1 and FOXB1 also play important roles in neural development (46-48). Behavioral misregulation is a key trait of ASD, and gene networks related to the GO term ‘behavior’ exhibited dysregulated expression in both affected individuals. Genes in these networks include *COMT, ADCYAP1, CNR1, HTR2C, GRIK2*, and *RGS4*, all of which are implicated in behavior-related phenotypes in humans and/or mice (49-57). ASD genes were also dysregulated in these affected individuals, relative to controls. Mutation of these genes in other individuals is implicated in autism risk or causation. These include adhesion-related genes (*PCDHA1*, *PCHDHA6*, *PCDHGA11, PCDH8, PCDH9, PCDH10* (58-62)*, KIRREL3* (63), *CNTN3*, *CNTN*4 (64), *CNTNAP4* (65), and *THBS1* (66)), receptor and channel genes (*CACNA2D3* and *SCN9A* (67)*, GRIK2* and *GRIK3* (68-71), *KCNJ2* (72, 73), and *GRIA1* (74, 75)), and genes associated with central nervous system development and axon guidance (*ERBB4* (76, 77), *NTNG1* (78), *TSHZ3* (79), *EBF3* (80)*, MYT1L* (81, 82), and *ANXA1* (40-42)). Altered expression of ASD-associated genes has also been observed in cellular models derived from affected individuals in other studies (8, 13-16, 19, 83). Therefore, these findings suggest that misregulated expression of suites of ASD-associated genes may contribute to risk or affectation, and may do so by altering neurodevelopment and/or neuronal function in these affected individuals.

A unique aspect of this study is the use of iPSC-based directed differentiation into both cExNs and cINs, enabling us to identify neural cell type-specific alterations associated with affectation. Although DEGs identified in affected individuals in both neural cell types were associated with many similar functions and diseases (e.g. behavior), the specific DEGs obtained often varied by cell type. For example, cIN DEGs included many more ASD-associated genes and protocadherin genes, the latter of which control neuronal migration, axonal growth, and synapse formation (60, 61). Human post-mortem cortical tissue from individuals with ASD has been shown to exhibit disrupted expression of cIN-associated genes, evidence that this cell type may commonly be disrupted in affected individuals *in vivo* (84). These findings suggest that extending cellular modeling studies to multiple disease-relevant neuronal cell types, including cINs, may reveal additional neurodevelopmental disruptions related to affectation.

To define cellular and molecular perturbations commonly related to affectation, we compared our findings to other studies that modeled ASD by directed differentiation of iPSCs into cExNs. We identified subsets of overlapping DEGs in comparisons with studies involving idiopathic autism cases vs. controls (26 shared DEGs; (7)), syndromic ASD involving macrocephaly (31 shared DEGs; (19)), and modeling of the syndromic ASD gene *CHD8 (*32 shared DEGs; (13)) (Additional file 6: Table S5). Data for such comparisons is limited at present because iPSC-based models have been generated for a relatively small number of individuals and mutations, and these almost exclusively characterize cExNs or cerebral organoids (7, 13, 18, 19).

The multiplex pedigree studied here was subjected to clinical exome sequencing, as it was hypothesized that a single, shared, genetic contributor might mediate autism risk and differential phenotypic expression by sex in this family. In this sequencing analysis, a thread of shared genetic liability amongst all children was a VUS in the ASD and ID-associated gene, *GPD2* (85-87), which was inherited from their mother. However, there is variable ASD expressivity amongst these individuals, ranging from absent, to intermediate, to severe. In addition, both males and females in the pedigree are variably affected, indicating the presence of other significant contributors to variation in severity of affectation within this family. This observation is consistent with recent evidence that genetic liability for ASD is prevalently polygenic, and that, even in multiplex pedigrees where a significant monogenic contributor has been identified, additional polygenic risk can contribute to affectation (37). Moreover, this multiplex family was prototypic in reflecting the most severe form of affectation occurring in a male.

We hypothesized that it might be possible to identify graded cellular phenotypes that correlated with the level of severity of phenotypic expression. In general, we instead observed many cellular and molecular alterations that were shared by the cellular models derived from the affected individuals, while not being observed in those derived from the unaffected individuals. However, we did define some proband-specific DEGs, not present in the less severely affected sister, many of which relate to behavior and nervous system development. A subset of these DEGs had graded expression, exhibiting intermediate expression levels in the intermediate phenotype sister, between her unaffected mother and her severely affected brother. These findings suggest that both the degree of dysregulation of expression and the number and identity of DEGs within these networks may contribute to the level of affectation. While further experimentation might reveal additional graded phenotypes, particularly in mature neurons, *ex vivo* cellular modeling cannot recapitulate many aspects of fetal and post-natal neurodevelopment that may have been perturbed to contribute to the graded affectation observed in these individuals.

This work highlights several considerations for ongoing scientific efforts to model this complex but prevalent form of ASD in future studies. First, since the unique characteristics of any multiplex pedigree present challenges for cellular modeling, it is important to control for sex and variation in affectation in subject and family selection, study design, and analysis. Related to this point is the importance of modeling affected females in such studies. Most ASD cellular modeling to date has been restricted to affected males (7-10, 13, 14, 16, 18, 19), given the increased prevalence of ASD among males, and the fact that constraint to a single sex simplifies some modeling considerations. In particular, sex chromosome dosage effects do not need to be accounted for in male cells, while female-derived iPSC models cannot currently recapitulate the process of random X-chromosome inactivation that occurs in developing somatic tissues, including the brain (88, 89). Interestingly, the transcriptomic differences that we observed here were not driven by sex-linked gene expression: very few DEGs in any potential pairwise sample comparison (whether between same or opposite sex models) were sex chromosome-linked potential contributors to sex-biased gene expression in the human brain (90, 91). Therefore, this work supports the feasibility of identifying DEGs associated with affectation by cellular model cross-comparisons, even when these models are derived from both female and male subjects.

Another consideration for iPSC-based modeling of ASD is genetic background, which can be a confounding variable for cross-comparisons (35). In this pedigree, ASD risk was polygenic, such that it was not possible to engineer a correction of a single genome variant to create pairs of isogenic mutant versus wild-type iPSC lines with an identical genetic background for study. In such cases, modeling of first degree relatives may serve as the best control, and modeling of multiple related individuals with varying affectation provides additional opportunities for identifying potential contributors to these differences in affectation. Including unrelated controls and performing comparisons with other studies can further highlight which phenotypic and transcriptomic alterations track with affectation, even by comparison with models derived from individuals with an unrelated genetic background.

## Conclusions

In summary, this work used robust schemes for differentiation of cortical neurons from iPSCs to model cellular and molecular signatures associated with multiplex ASD in a family reflecting varying degrees of affectation. Even in this prevalent, complex form of ASD, involving heritability, polygenic etiology, and variable affectation, we could identify affectation-linked cellular and molecular alterations of neurodevelopment, some of which overlapped those defined in other iPSC-based studies of monogenic, syndromic, and *de novo* ASD. As more cellular models of ASD are characterized, these data can be harnessed in the search for convergent and divergent contributors to impairment across the genetically complex and multi-factorial pathways that give rise to ASD.

## Abbreviations

ASD: Autism Spectrum Disorder
iPSCs: Induced Pluripotent Stem Cells
cExNs: Cortical Excitatory Neurons
cINs: Cortical Inhibitory Neurons
NPCs: Neural Progenitor Cells
ES: Exome Sequencing
UM: Unaffected Mother
AP: Affected Proband
IS: Intermediate Phenotype Sister
UC: Unaffected Control
GTAC: Genome Technology Access Center
ICC: Immunocytochemistry
RT-qPCR: quantitative reverse transcription PCR
EBs: Embryoid Bodies
DEG: Differentially Expressed Gene
IPA: Ingenuity Pathway Analysis
VUS: Variant of Unknown Significance
PI: Propidium Iodide
PCA: Principal Component Analysis
GO: Gene Ontology
SFARI: Simons Foundation Autism Research Initiative.

## Ethics approval and consent to participate

Subjects were consented for biobanking and iPSC line generation by the Washington University Institutional Review Board of the Human Research Protection Office under human studies protocol #201409091 (Dr. John Constantino).

## Consent for publication

Consent to publish data was provided by all subjects.

## Availability of data and materials

The RNA-seq data generated during the current study are available in the Gene Expression Omnibus (GEO) repository as Series GSE129806. The ASD gene dataset analyzed during the current study is available in the SFARI Gene database at https://gene.sfari.org.

## Author’s contributions

E.M.A.L. contributed to the study design, carried out all cExN experimentation, analyzed data, and prepared the manuscript. K.M. contributed to the study design, carried out all cIN experimentation, analyzed data and contributed to manuscript preparation. D.B. interpreted clinical exome sequencing data and contributed to manuscript preparation. P.G. and B.Z. performed RNA-seq data analysis. A.B. contributed to the study design. J.N.C. contributed to the study design and manuscript preparation. K.L.K. contributed to the study design, data analysis, and manuscript preparation. All authors read and approved the final manuscript.

## Acknowledgments

We thank the family for providing biomaterials for use in this study. We thank the Genome Engineering and iPSC Center (GEiC) at Washington University School of Medicine (WUSM) for deriving the iPSC lines used in this study. We thank the Alvin J. Siteman Cancer Center at WUSM, for the use of the Siteman Flow Cytometry Core, which provided self-service Flow Cytometry Analysis. The Siteman Cancer Center is supported in part by an NCI Cancer Center Support Grant #P30 CA091842. We thank the Genome Technology Access Center in the Department of Genetics at WUSM for providing genomic sequencing services. The Center is partially supported by NCI Cancer Center Support Grant #P30 CA91842 to the Siteman Cancer Center and by ICTS/CTSA Grant# UL1 TR000448 from NIH/NCRR.

## Additional files

**Additional file 1: Table S1.** Antibodies used in immunocytochemistry and immunoblotting experiments (XLSX).

**Additional file 2: Figures S1-S5.** Figure legends provided in file (PDF).

**Additional file 3: Table S2.** DEGs from pairwise comparisons of the four sample types for cExN or cIN NPCs (XLSX).

**Additional file 4: Table S3.** Hierarchical clustering of the relative expression of DEGs across the four sample types for cExN and cIN NPCs was performed using ClustVis. Order of genes in the cluster and unit variance scaled relative expression values are indicated (XLSX).

**Additional file 5: Table S4.** Details about selected DEGs within IPA networks (PDF).

**Additional file 6: Table S5.** cExN DEG comparison to other studies (XLSX).

