## Additional File 2 (pdf) for "Cellular and molecular characterization of multiplex autism in human induced pluripotent stem cell-derived neurons"

**Additional file 2: Figures S1-S5.**

**Supplemental Figure S1. Characterization of iPSCs. (A)** Exome sequencing identified a heterozygous missense variant in the FAD domain-encoding region of the *GPD2* gene (chr2: 157352686 (hg19) G>A, NM_001083112.2 c.233G>A, p.G78E) in individuals in the pedigree under study, including the unaffected mother (UM), intermediate phenotype sister (IS) and affected proband (AP), as well as in the trait-affected brother (TB) not studied here. **(B)** This variant was confirmed by Sanger sequencing and was not present in an unrelated, unaffected control (UC). **(C-E)** Expression of markers of pluripotency were assessed by (C) RT-qPCR and (D) immunocytochemistry (representative images shown), comparing the cell lines under study (n=3 biological replicates from one clonal line for each subject). GPD2 protein level was also assessed in these iPSC lines by (D) immunocytochemistry or (E) Western blotting (n=2 biological replicates from one clonal line for each subject). **(F-G)** iPSCs were stained with propidium iodide and analyzed by FACS for DNA content analysis. (F) Representative FACS plots are shown. (G) Mean percentages of cells in each cell cycle stage, with no differences between cell lines observed (n=3 biological replicates from one clonal line for each subject). p-values: *$P$<0.05, **$P$<0.01, ***$P$<0.001 were determined by a two-tailed Student's t-test and all other pairwise comparisons had a non-significant p-value ($P$≥0.05). In D, scale bar = 200µm.

**Supplemental Figure S2. Cellular characterization of iPSC-derived cExN and cIN NPCs. (A-B)** After 12 days of differentiation, cExN NPCs were stained with propidium iodide and analyzed by FACS for DNA content. (A) Representative FACS plots. (B) Mean percentages of cells in each cell cycle stage, with no differences visible between cell lines (n=4 biological replicates from one clonal line for each subject). **(C-D)** cExN NPCs were plated in equal numbers for each sample and were counted after four days of culture (n=6 biological replicates from one clonal line for each subject). Data are quantified in (C), and representative images are shown in (D). **(E)** GPD2 protein levels were detected in cExN NPC samples by Western blotting (n=2 biological replicates from one clonal line for each subject). **(F)** After 12 days of differentiation, cIN NPCs were stained with propidium iodide and analyzed by FACS for DNA content. Mean percentages of cells in each cell cycle stage are shown (n=3 biological replicates). **(G-H)** cIN NPCs were plated in equal numbers for each sample and counted after four days of culture. Data are quantified in (G) and representative images are shown in (H) (n=3 biological replicates from one clonal line for each subject). p-values: *$P$<0.05, **$P$<0.01, ***$P$<0.001 were determined by a two-tailed Student's t-test and all other pairwise comparisons had a non-significant p-value ($P$≥0.05). In D and H scale bar = 300µm.

**Supplemental Figure S3. Maturation of cExN NPCs. (A)** Differentiation scheme, including timeline and small molecules used. **(B)** Immunocytochemistry for proteins marking early cExN specification, maturation, and neuronal function (scale bar = 50μm) (n=1 biological replicate from one clonal line for each subject; representative images are shown).

**Supplemental Figure S4. Transcriptome analysis of cExN and cIN NPCs by RNA-seq. (A-B)** Multidimensional scaling plots for the (A) cExN NPC samples and the (B) cIN NPC samples (B). **(C-D)** Numbers of differentially expressed genes obtained for each pairwise sample comparison, indicating up- and down-regulated genes for the (C) cExN NPCs and (D) cIN NPCs ($p<0.05$, fold-change >2).

**Supplemental Figure S5**. **Networks of DEGs with similar expression in the AP/IS and different than the UM.** Networks of DEGs are related to **(A)** cExN **'**neurological disease' **(B)** cIN **'**nervous system development and function', and **(C)** cIN **'**neurological'. Within each network, red symbols indicate upregulated genes and green symbols indicate downregulated genes, where the color intensity represents relative degree of differential expression.

**S. Fig. S1**

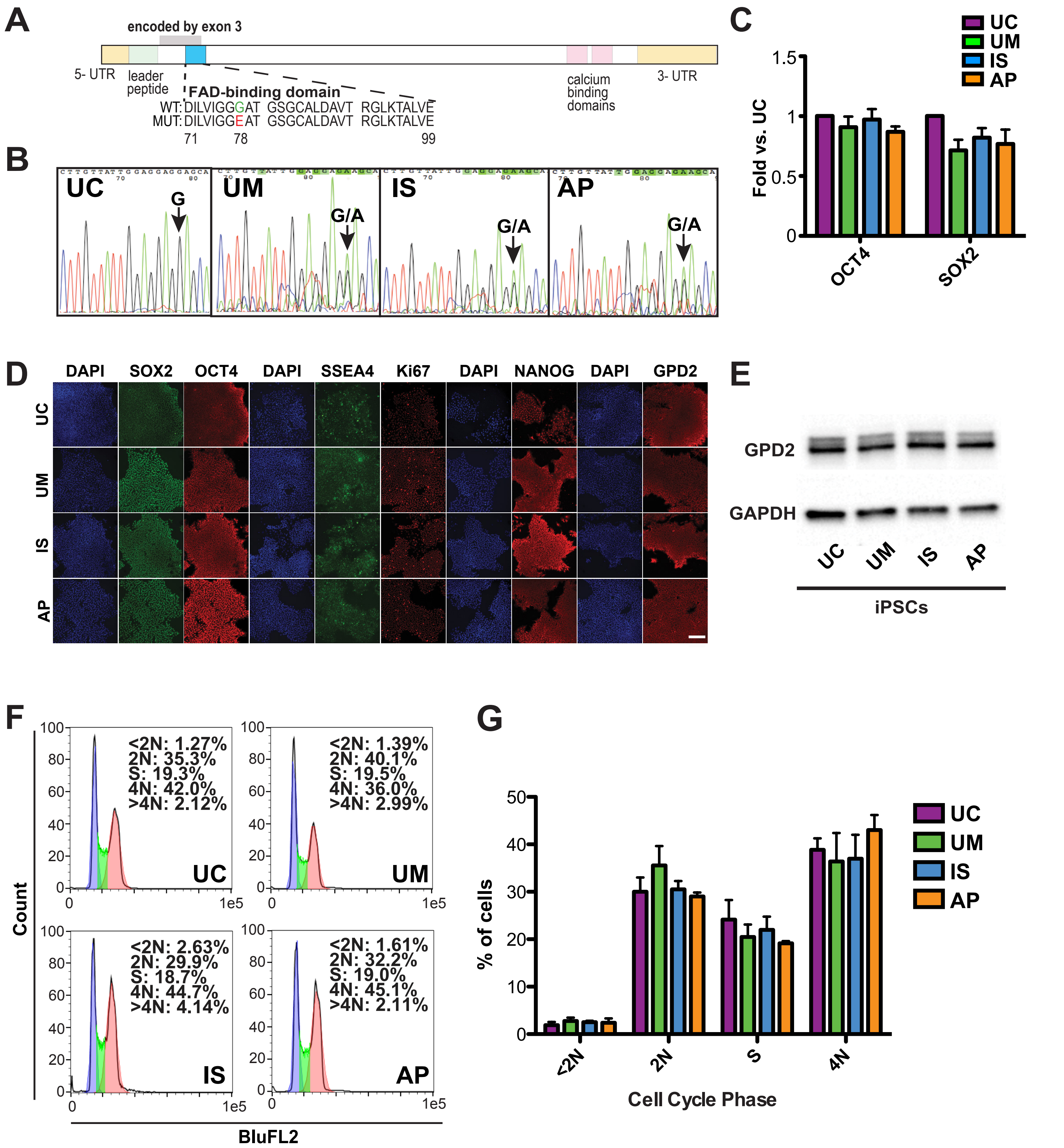
A
encoded by exon 3
5- UTR
leader peptide
FAD-binding domain
WT: DILVIGGGAT GSGCALDAVT RGLKTALVE
MUT: DILVIGGEAT GSGCALDAVT RGLKTALVE
71
78
99
calcium binding domains
3- UTR
B
UC
G
UM
G/A
IS
G/A
AP
G/A
C
UC
UM
IS
AP
Fold vs. UC
1.5
1
0.5
0
OCT4
SOX2
D
DAPI
SOX2
OCT4
DAPI
SSEA4
Ki67
DAPI
NANOG
DAPI
GPD2
UC
UM
IS
AP
E
GPD2
GAPDH
UC
UM
IS
AP
iPSCs
F
Count
<2N: 1.27%
2N: 35.3%
S: 19.3%
4N: 42.0%
>4N: 2.12%
UC
<2N: 1.39%
2N: 40.1%
S: 19.5%
4N: 36.0%
>4N: 2.99%
UM
<2N: 2.63%
2N: 29.9%
S: 18.7%
4N: 44.7%
>4N: 4.14%
IS
<2N: 1.61%
2N: 32.2%
S: 19.0%
4N: 45.1%
>4N: 2.11%
AP
100
80
60
40
20
0
0
1e5
BluFL2
G
% of cells
50
40
30
20
10
0
<2N
2N
S
4N
Cell Cycle Phase
UC
UM
IS
AP

### S. Fig. S2

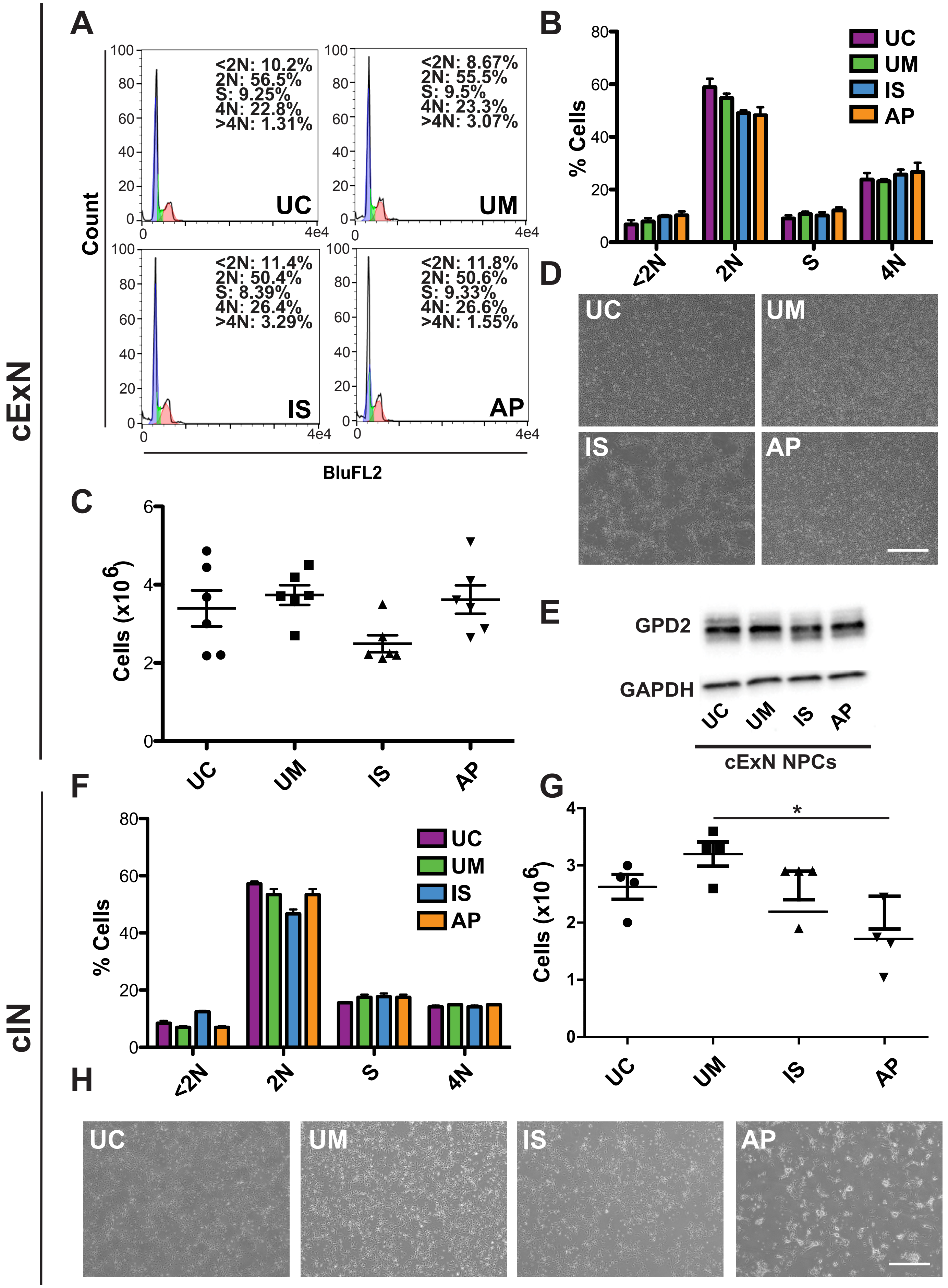
cExN
A
<2N: 10.2%
2N: 56.5%
S: 9.25%
4N: 22.8%
>4N: 1.31%
UC
<2N: 8.67%
2N: 55.5%
S: 9.5%
4N: 23.3%
>4N: 3.07%
UM
<2N: 11.4%
2N: 50.4%
S: 8.39%
4N: 26.4%
>4N: 3.29%
IS
<2N: 11.8%
2N: 50.6%
S: 9.33%
4N: 26.6%
>4N: 1.55%
AP
Count
BluFL2
B
% Cells
UC
UM
IS
AP
<2N
2N
S
4N
C
Cells (x10^6)
UC
UM
IS
AP
D
UC
UM
IS
AP
E
GPD2
GAPDH
UC
UM
IS
AP
cExN NPCs
cIN
F
% Cells
UC
UM
IS
AP
<2N
2N
S
4N
G
Cells (x10^6)
*
UC
UM
IS
AP
H
UC
UM
IS
AP

S. Fig. S3

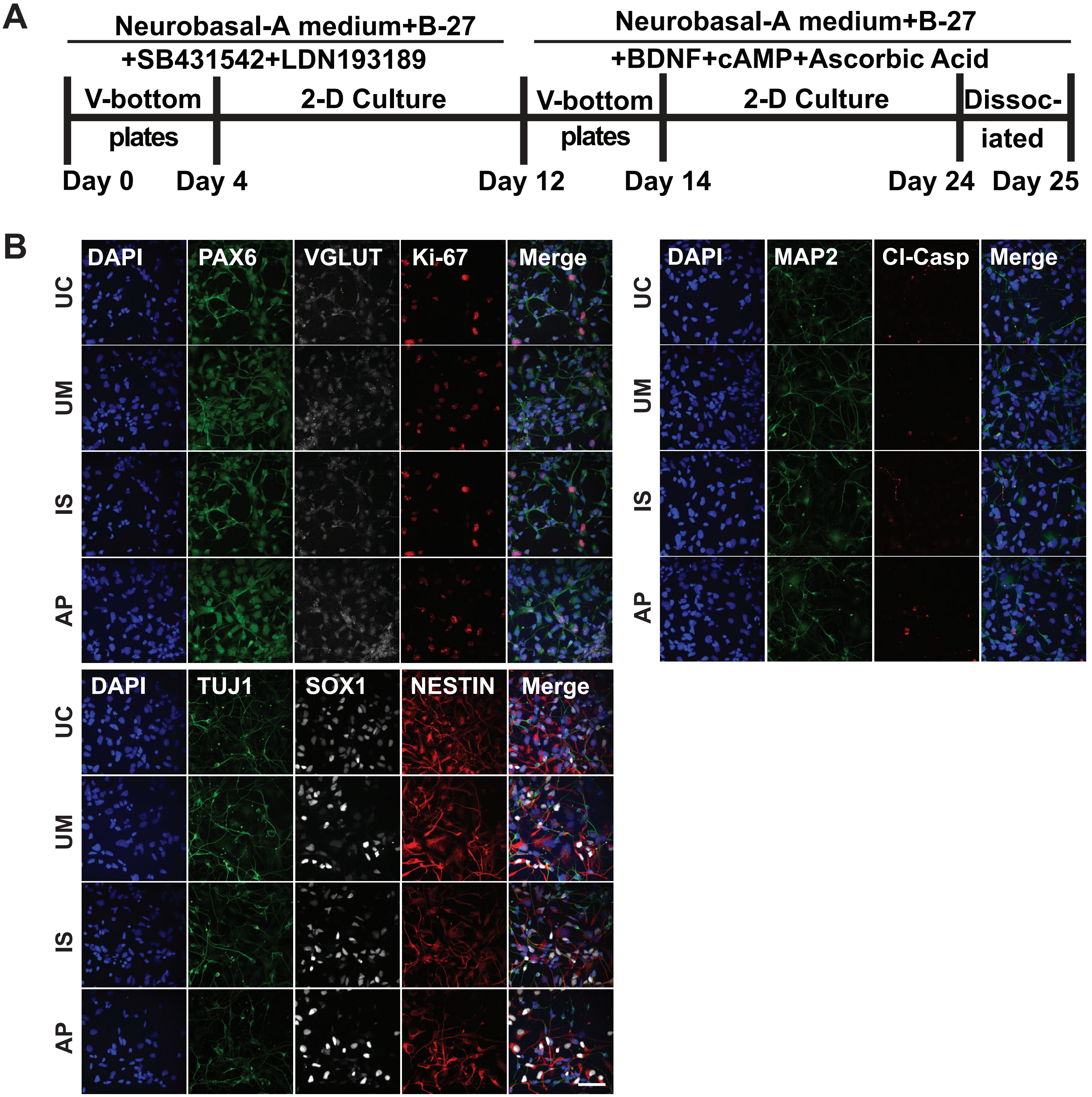
A
Neurobasal-A medium+B-27
+SB431542+LDN193189
Neurobasal-A medium+B-27
+BDNF+cAMP+Ascorbic Acid
V-bottom plates
2-D Culture
V-bottom plates
2-D Culture
Dissoc-iated
Day 0
Day 4
Day 12
Day 14
Day 24
Day 25
B
DAPI
PAX6
VGLUT
Ki-67
Merge
UC
UM
IS
AP
DAPI
MAP2
Cl-Casp
Merge
DAPI
TUJ1
SOX1
NESTIN
Merge

**S. Fig. S4**

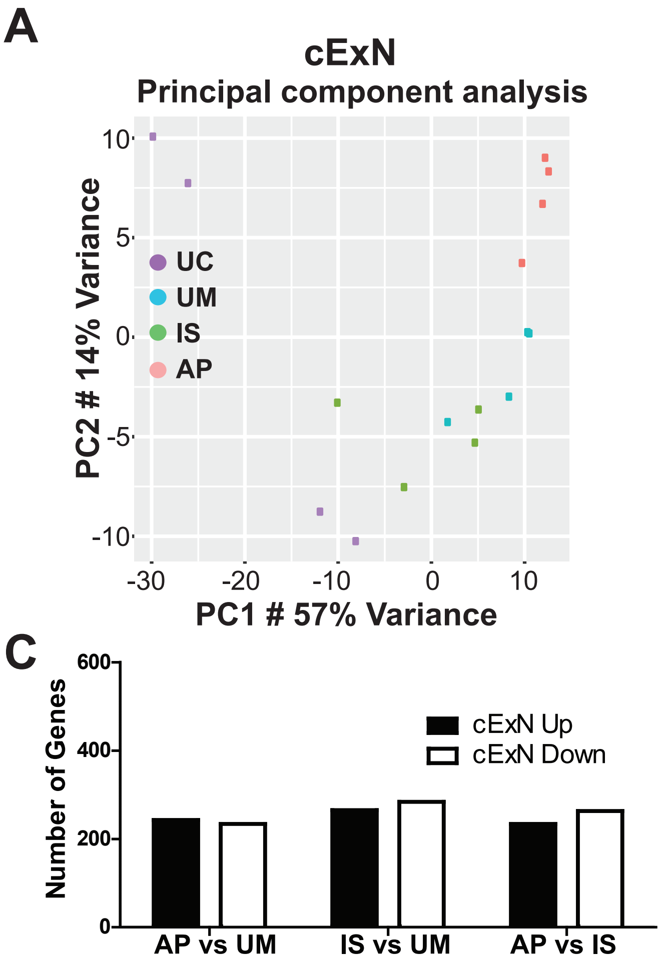
A
cExN
Principal component analysis
PC2 # 14% Variance
PC1 # 57% Variance
UC
UM
IS
AP
10
5
0
-5
-10
-30
-20
-10
0
10
B
cIN
Principal component analysis
PC2 # 30% Variance
PC1 # 39% Variance
UC
UM
IS
AP
5
0
-5
-10
-15
-10
-5
0
5
10
C
Number of Genes
600
400
200
0
cExN Up
cExN Down
AP vs UM
IS vs UM
AP vs IS
D
Number of Genes
600
400
200
0
cIN Up
cIN Down
AP vs UM
IS vs UM
AP vs IS

**S. Fig. S5**

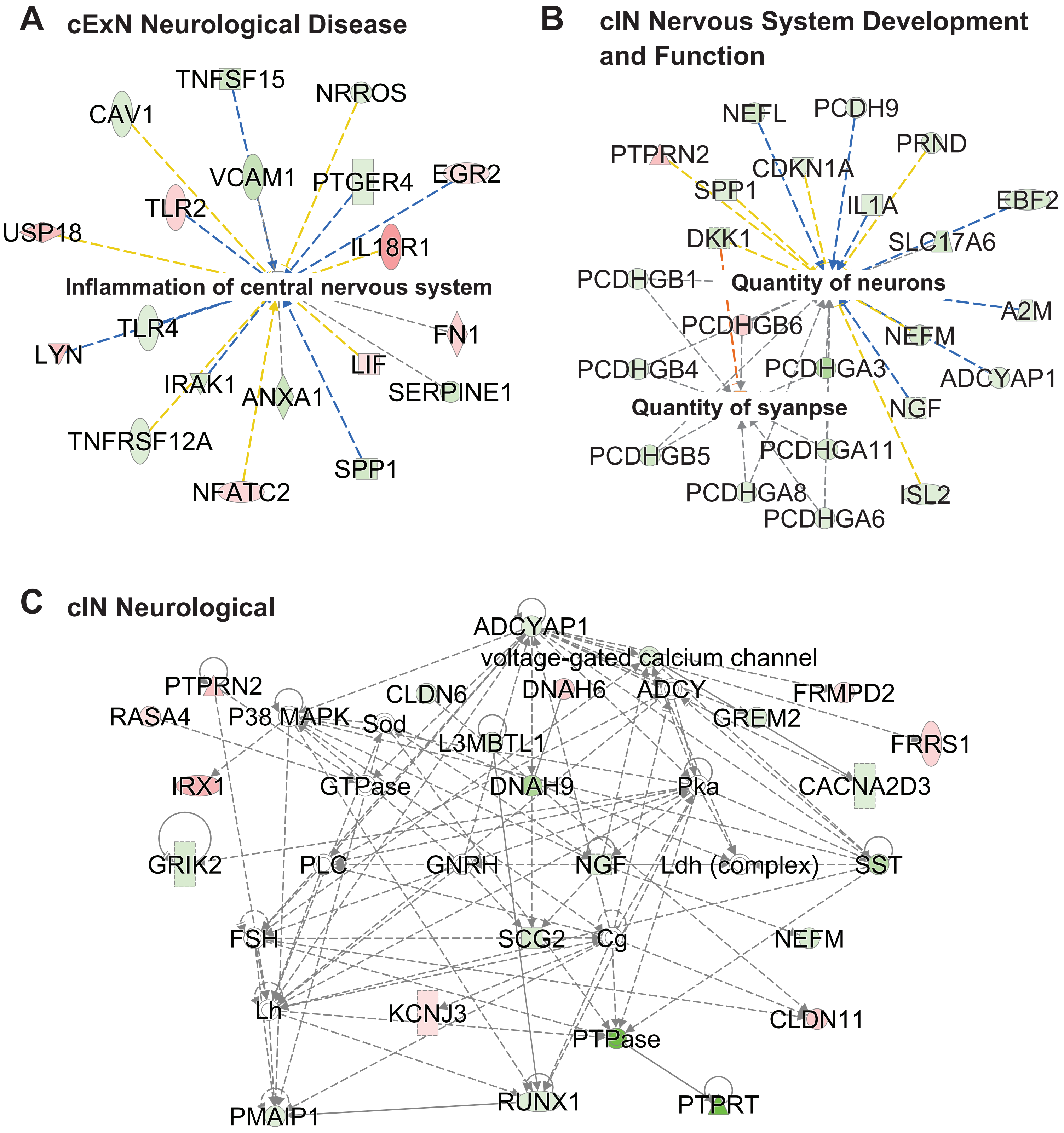
A
cExN Neurological Disease
TNFSF15
NRROS
CAV1
VCAM1
PTGER4
EGR2
TLR2
USP18
IL18R1
Inflammation of central nervous system
TLR4
LYN
FN1
LIF
IRAK1
ANXA1
SERPINE1
TNFRSF12A
SPP1
NFATC2
B
cIN Nervous System Development and Function
NEFL
PCDH9
PRND
PTPRN2
CDKN1A
SPP1
IL1A
EBF2
DKK1
SLC17A6
PCDHGB1
Quantity of neurons
A2M
PCDHGB6
NEFM
PCDHGB4
PCDHGA3
ADCYAP1
Quantity of syanpse
NGF
PCDHGB5
PCDHGA11
PCDHGA8
ISL2
PCDHGA6
C
cIN Neurological
ADCYAP1
voltage-gated calcium channel
PTPRN2
CLDN6
DNAH6
ADCY
FRMPD2
RASA4
P38 MAPK
Sod
GREM2
L3MBTL1
FRRS1
IRX1
GTPase
DNAH9
Pka
CACNA2D3
GRIK2
PLC
GNRH
NGF
Ldh (complex)
SST
FSH
SCG2
Cg
NEFM
Lh
KCNJ3
CLDN11
PTPase
PMAIP1
RUNX1
PTPRT
