## Additional File 5 (pdf) for "Cellular and molecular characterization of multiplex autism in human induced pluripotent stem cell-derived neurons"

**Additional file 5 – Table S4 – Details about selected DEGs within IPA networks.**

| cExN or cIN, IS/AP shared or AP-specific | Gene Name | Class | Network identified | Physiological role and function | References |
|---|---|---|---|---|---|
| cExN – IS/AP shared | *CHL1* | Adhesion molecule | Behavior (Locomotion), Behavior and Developmental Disorder | Nervous system development and synaptic plasticity. | [25, 26] |
| | *KCNC1* | Ion channel | Behavior (Locomotion) | Potassium channel, loss associated with epilepsy and ID. | [27] |
| | *SPP1* | ECM protein | Behavior (Locomotion), Behavior and Developmental Disorder, Neurological Disease (Inflammation of Central Nervous System) | Neuroprotective, enhances NSC survival and proliferation. | [28, 29] |
| | *TPH1* | Enzyme | Behavior (Locomotion), Behavior and Developmental Disorder | Serotonin biosynthesis, mutated in schizophrenia and other neuropsychiatric disorders. | [30, 31] |
| | *GAS7* | PCH protein | Behavior and Developmental Disorder | Adaptor protein; regulate cytoskeleton/membrane dynamics. Involved in neurite outgrowth. | [32, 33] |
| | *XYLT1* | Xylosyl transferase | Behavior and Developmental Disorder | Involved in proteoglycan synthesis, which has a role in neuronal migration. | [34, 35] |
| | *SPATA18* | Mitochondria-eating protein | Behavior and Developmental Disorder | Mitochondrial response to oxidative stress. | [36] |
| | *CHCHD2* | Mitochondrial protein | Behavior and Developmental Disorder | Regulates metabolism and scavenging reactive oxygen species. Negative regulator of mitochondria-mediated apoptosis. | [37] |
| | *VCAM1* | Adhesion | Neurological Disease (Inflammation of Central Nervous System) | Maintains NSC identity and adult NSC niche. | [38, 39] |
| | *ANXA1* | Annexin | Neurological Disease (Inflammation of Central Nervous System) | Proliferation, differentiation, apoptosis. Anti-inflammatory. Recurrent duplications associated with ASD. | [40-42] |
| | *SERPINE1* | Serine proteinase inhibitor | Neurological Disease (Inflammation of Central Nervous System) | Part of MET signaling cascade, which has been associated with ASD. Role in | [43, 44] |

| | | | | | |
|---|---|---|---|---|---|
| | | | | brain not known. Upregulated in human NSCs versus other brain tissue. | |
| | *TLR4 and TLR2* | Toll-like receptor | Neurological Disease (Inflammation of Central Nervous System) and Behavior (Locomotion) | Neuronal differentiation and survival. | [45, 46] |
| | *IRAK1* | Kinase – member of Toll/IL-1-receptor family | Neurological Disease (Inflammation of Central Nervous System) | Might contribute to neuroprotection. | [47] |
| cExN – AP-specific | *ERBB4* | EGF receptor tyrosine kinase | Behavior (Memory and Learning) and Nervous System Development and Function (Differentiation of Neurons) | Proliferation, differentiation, migration, and survival of neural cells. | [48, 49] |
| | *FOXB1* | Transcription factor | Behavior (Memory and Learning) | Expressed in neural tube, involved in anterior-posterior patterning and in neural development during embryogenesis | [50, 51] |
| | *COMT* | Catechol-O-methyltransferase | Behavior (Memory and Learning) | Breaks down dopamine to maintain normal physiological levels in the prefrontal cortex. | [52] |
| | *SLC8A2* | Sodium/ calcium exchanger | Behavior (Memory and Learning) | Involved in synaptic plasticity. | [53] |
| | *EMX1* | Transcription factor | Nervous System Development and Function (Differentiation of Neurons) | Central role in neural development. | [54] |
| cIN – IS/AP shared | *KCNJ3* and *KCNJ2* | Ion channels | Behavior (Behavior), Neurological | Behavior, mood disorder, and motor coordination. | [1-4] |
| | *CACNA2D3* | Ion channel | Behavior (Behavior), Neurological, Psychological Disorder (Anxiety Disorders) | Mood and cognition. | [5] |
| | *SCN9A* | Ion channel | Psychological Disorder (Anxiety Disorders and Depressive Disorder) | Excitability of sensory and cortical neurons. | [6] |

| | | | | | |
|---|---|---|---|---|---|
| | *ADCYAP1* | Neuropeptide | Behavior (Learning, Cognition, and Behavior), Nervous System Development and Function (Quantity of Neurons) | Regulation of psychomotor and sensory motor behavior and social interactions. | [7, 8] |
| | *GRIK2, GRIK3* | Receptors | Psychological Disorder (Mood Disorders), Behavior (Behavior), Neurological | Motor activity and habituation. | [1, 9, 10] |
| | *SST* | Calcium binding protein | Behavior (Learning, Cognition, and Behavior), Neurological | Mood disturbances. | [11] |
| | *PCDH9* and *PCDHGA11* | Adhesion molecules | Behavior (Learning, Cognition, and Behavior), Nervous System Development and Function (Quantity of Synapse and Quantity of Neurons) | Learning, memory, behavior, neuronal migration, axonal growth, and synaptic function. | [12-16] |
| | *SYT4* | Transcription factor | Behavior (Learning and Cognition) | Synaptic transmission and mental retardation. | [17, 18] |
| cIN - AP-specific | *GRIA1 and GRIA2* | Receptors | Behavior (Behavior) and Nervous System Development and Function (Synaptic Transmission) | Synaptic structural and functional plasticity. | [19] |
| | *GAP43* | Gap junction | Nervous System Development and Function (Development of Neurons) | Stress and abnormal behavior. | [20] |
| | *ARC* | Cytoskeleton protein | Nervous System Development and Function (Synaptic Transmission), Behavior (Behavior and Cognition) | Synaptic plasticity and memory. | [21] |
| | *MYT1L* | Transcription factor | Nervous System Development and Function (Development of Neurons) | Brain development and intellectual disability. | [22, 23] |
| | *CNTN1 and CNTN2* | Adhesion molecules | Behavior (Behavior and Cognition), Nervous System Development (Development of Neurons) | Nervous system development. | [24] |
